# Derlin rhomboid pseudoproteases employ substrate engagement and lipid distortion function for retrotranslocation of ER multi-spanning membrane substrates

**DOI:** 10.1101/2021.06.15.448428

**Authors:** Anahita Nejatfard, Nicholas Wauer, Satarupa Bhaduri, Adam Conn, Saroj Gourkanti, Narinderbir Singh, Tiffany Kuo, Rachel Kandel, Rommie E. Amaro, Sonya E. Neal

## Abstract

Nearly one-third of proteins are initially targeted to the endoplasmic reticulum (ER) membrane where they are correctly folded, assembled, and then delivered to their final cellular destinations. In order to prevent the accumulation of misfolded membrane proteins, ER associated degradation (ERAD) moves these clients from the ER membrane to the cytosol; a process known as retrotranslocation. Our recent work in *S. cerevisiae* has revealed a derlin rhomboid pseudoprotease, Dfm1, is involved in the retrotranslocation of ubiquitinated ERAD membrane substrates. In this study, we sought to understand the mechanism associated with Dfm1’s actions and found that Dfm1’s conserved rhomboid residues are critical for membrane protein retrotranslocation. Specifically, we identified several retrotranslocation-deficient Loop 1 mutants that display impaired binding to membrane substrates. Furthermore, Dfm1 has retained the lipid thinning functions of its rhomboid protease predecessors to facilitate in the removal of ER membrane substrates. We find this substrate engagement and lipid thinning feature is conserved in its human homolog, Derlin-1. Utilizing interaction studies and molecular dynamics simulations, this work reveals that derlin rhomboid pseudoproteases employ novel mechanisms of substrate engagement and lipid thinning for catalyzing extraction of multi-spanning membrane substrates.

## Introduction

Almost all eukaryotic membrane and secreted proteins, comprising one third of the eukaryotic proteome, are co-translationally imported into the endoplasmic reticulum (ER) and where they are subsequently folded(Sicari et al., 2019; Wang and Dehesh, 2018). Often, proteins fail to fold or assemble properly, at which point they are eliminated by ER-Associated-Degradation (ERAD)(Mehrtash and Hochstrasser, 2019a; Ruggiano et al., 2014; Sun and Brodsky, 2019). ERAD is a highly conserved quality control pathway that selectively targets misfolded proteins for degradation. Defective ERAD results in a toxic buildup of damaged proteins associated with many maladies such as aging, neurodegenerative diseases, retinal degeneration, diabetes, rheumatoid arthritis (Bhattacharya and Qi, 2019; Xiong et al., 2020; Yagishita et al., 2008). Furthermore, many pathogens hijack ERAD machinery to deliver toxins to their cytosolic targets (Cho et al., 2012). ERAD has both deleterious and advantageous roles in cancer progression; holding the promise for therapeutic use upon better understanding(Moon et al., 2018). As a result, ERAD has multiple impacts on normal physiology and human disease.

ERAD involves several key steps(Hampton and Garza, 2009; Mehrtash and Hochstrasser, 2019): **1)** recognition of misfolded ER proteins, **2)** polyubiquitination of substrates by one or more E3 ligases, **3)** movement or extraction of ER substrates to the cytosol (termed retrotranslocation), which is powered by Cdc48/p97 AAA-ATPase, and **4)** degradation by the cytosolic proteasome. ERAD recognizes different classes of substrates based on the location of the lesion within the protein. Substrates for ERAD include misfolded soluble luminal proteins (ERAD-L) and integral membrane proteins with lesions in their transmembrane domain (ERAD-M)(Hampton et al., 1996; Plemper et al., 1998; Vashist and Ng, 2004; Wangeline and Hampton, 2018) or their cytosolic domain (ERAD-C)(Ravid et al., 2006). The HRD pathway uses E3 ubiquitin ligase Hrd1 for ubiquitination of both ERAD-M and ERAD-L substrates, whereas the DOA (Degradation of alpha 2) pathway uses E3 ubiquitin ligase Doa10 for ubiquitination of ERAD-C substrates (Carvalho et al., 2006; Chen et al., 2006; Foresti et al., 2013; Hampton and Garza, 2009; Laney and Hochstrasser, 2003).

A common feature of all ERAD pathways is the requirement for moving misfolded substrates from the ER to the cytosol for degradation; a process known as retrotranslocation (Hampton and Sommer, 2012). Derlins and E3 ligases have been implicated in removing substrates from the ER(Greenblatt et al., 2011; Lilley and Ploegh, 2004; Ye et al., 2004). A recent study has shown that yeast derlin, Der1, along with E3 ligase Hrd1 forms a channel to assist in the retrotranslocation of ER luminal substrates (Wu et al., 2020). Furthermore, E3 ligase Doa10 was demonstrated to serve as a retrotranslocon for a single-spanning tail-anchored substrate(Peterson et al., 2019; Schmidt et al., 2020; Wu et al., 2020). In contrast, the transmembrane domains of Hrd1 and Doa10 were not required for retrotranslocating a subset of ERAD membrane substrates (Garza et al., 2009a). Despite intense study on this pathway, the protein required for retrotranslocating multi-spanning membrane substrates remained unknown for over two decades and has only been recently identified. We have previously explored membrane substrate retrotranslocation by screening a complete collection of yeast mutants via SPOCK (single-plate orf compendium kit), which consists of a 5,808 yeast strain array of non-essential gene deletion mutants and essential DAmP gene mutants, for deficiencies in degradation of SUS (self-ubiquitinating substrate)-GFP. These studies pinpointed yeast derlin, Dfm1 as a specific mediator for the retrotranslocation of multi-spanning membrane substrates in ERAD (Neal et al., 2018). This finding contradicted previous results in which Dfm1 had either a partial or no role in ERAD (Avci et al., 2014; Goder et al., 2008; Sato and Hampton, 2006; Stolz et al., 2010). We resolved this discrepancy through the finding that loss of Dfm1 along with strong expression of membrane substrates imposes a growth stress on cells and causes rapid and complete suppression of the *dfm1Δ* retrotranslocation defect(Neal et al., 2018, 2020). Accordingly, the fast generation of test strains inherent in the SPOCK screen has revealed yeast derlin Dfm1 as a major mediator for membrane substrate retrotranslocation.

Sequence and structural homology approaches indicate that derlins belong to the rhomboid superfamily(Greenblatt et al., 2011). The rhomboid superfamily is known for their many roles in diverse membrane-related processes such as development, signaling, parasitic invasion, and protein trafficking, and protein quality control(Düsterhöft et al., 2017; Kandel and Neal, 2020). Many rhomboid proteins are intermembrane proteases and typically cleave membrane substrates within the lipid bilayer via the serine-histidine dyad active site (Bondar et al., 2009; Lemieux et al., 2007; Shokhen and Albeck, 2017; Tichá et al., 2018; Uritsky et al., 2016; Zhou et al., 2012). However, there is a large subclass of the rhomboid superfamily, to which derlins belong, that lack residues for proteolysis and are known as rhomboid pseudoproteases (Began et al., 2020; Lemberg and Adrain, 2016; Lemberg and Freeman, 2007). Despite the absence of the proteolytic site, rhomboid pseudoproteases have conserved rhomboid residues, implying the intriguing idea that derlins utilize rhomboid features for executing their retrotranslocation function. In support of this idea, we and others previously published that both human and yeast derlins require the conserved rhomboid motifs,WR and GxxxG, for substrate removal across the membrane(Greenblatt et al., 2011; Neal et al., 2018).

The structures of *E. coli* and *H. influenzae* rhomboid protease GlpG (Brooks and Lemieux, 2013; Lemieux et al., 2007; Wang et al., 2006) and an array of mechanistic studies on the rhomboid superfamily have elucidated some of the mechanistic principles that may be at play in derlin-mediated retrotranslocation. Bacterial rhomboids’ compact architectural fold is presumed to induce local perturbations of the lipid bilayer for gaining substrate access prior to cleavage (Bondar et al., 2009; Moin and Urban, 2012). An intriguing possibility is that Dfm1 has retained membrane perturbing properties of its bacterial counterpart to facilitate the movement of substrates across the membrane. In support of this idea, Dfm1’s yeast paralog, Der1, has been shown to induce lipid thinning to assist in the retrotranslocation of ER luminal substrates (Wu et al., 2020). However, it is yet to be determined whether the same mechanism is applied for the removal of integral membrane proteins from the ER. Accordingly, salient questions arising from these studies are: how does Dfm1 retrotranslocate ER membrane substrates, and does Dfm1 employ the same lipid distortion function for retrotranslocation?

In the studies below, we have explored these questions by performing a non-biased sequence analysis of Dfm1 coupled with cell biological assays and computational simulation to characterize the mechanistic features associated with retrotranslocation. Herein, we identified a subset of retrotranslocation-deficient mutants that are enriched in Loop 1 (L1) and transmembrane 2 (TM2) regions of Dfm1. Closer analysis reveals L1 retrotranslocation-deficient mutants are unable to bind to ERAD membrane substrates, indicating a role for L1 region in substrate detection. Furthermore, molecular dynamic simulations on Dfm1 homology model, revealed that Dfm1 possesses membrane lipid thinning function. Indeed, a retrotranslocation-deficient mutants are localized at TM2, which is critical for lipid thinning as delineated by computational modeling. Notably, Dfm1 retrotranslocation-deficient mutants are located at sites that are highly conserved in mammalian derlins. We sought to determine the conservation of Dfm1’s mechanism and found that a subset of Dfm1 sites identified from our screen also contribute to the membrane substrate binding and the lipid thinning function of human derlin, Derlin-1. Derlin-1 has been previously shown to utilize its rhomboid features for retrotranslocation and is implicated in several pathologies including cancer, cystic fibrosis, neuropathies and viral infection (Greenblatt et al., 2011; Kandel and Neal, 2020). Overall, our study sheds light on how derlin rhomboid pseudoproteases have evolved to carry out the critical and widely conserved process of membrane protein quality control.

## Results

### Yeast Dfm1 has highly conserved rhomboid and derlin-specific residues

Sequence alignment of Dfm1 along with other members of the rhomboid superfamily reveals Dfm1 contains residues that are highly conserved across the rhomboid superfamily (Fig. S1A; *highlighted in red and yellow*). Along with having common rhomboid features, Dfm1 has conserved residues that are specifically retained in mammalian derlin rhomboid pseudoproteases (Fig. S1A; *encircled in orange*). In this study, we sought to understand the extent that rhomboid and/or derlin-specific features of Dfm1 are required for membrane substrate retrotranslocation.

### Rhomboid WR and GxxxG is not sufficient for retrotranslocation

Dfm1 contains the highly conserved WR motif in L1 and a GxxxG motif in TM6. We and others have shown both motifs are important for rhomboid derlin retrotranslocation function (Greenblatt et al., 2011; Neal et al., 2018). We examined whether the WR and GxxxG motifs are sufficient for Dfm1 retrotranslocation function through use of chimeras. We used the Der1-SHP chimera from our previous studies, which consists of Dfm1’s closest homolog, Der1, fused to Dfm1’s cytoplasmic SHP tail (Fig. 1A)(Neal et al., 2018). Our previous work indicated Der1-SHP supports Cdc48 recruitment via binding of Cdc48 to the chimera’s SHP tail, but doesn’t support retrotranslocation through Der1’s transmembrane segment (Fig. 1A)(Neal et al., 2018). A closer examination of Der1’s transmembrane regions show Der1 does not possess the highly conserved WR and GxxxG motif and harbors **G**R and **N**xxxG instead (Fig. 1B). To determine whether the WR and GxxxG motifs are sufficient in supporting Der1-Shp retrotranslocation function, both motifs were inserted at homologous sites within the Der1-Shp transmembrane region. Three mutants were generated: Der1-Shp-**WR**, Der1-Shp-**GxxxG,** and Der1-Shp- **WR+GxxxG**. All three Der1-Shp mutants had the same stability as the unaltered Der1-Shp chimera protein and were still able to support Cdc48 recruitment as indicated with a direct assay of Cdc48 binding to the microsomal pellet (Fig. 1C,E). Despite this, neither mutant could facilitate degradation of Hmg2-GFP through ERAD, implying additional sequence residues within Dfm1’s transmembrane segments are required for retrotranslocation (Fig.1D).

**Figure 1.**
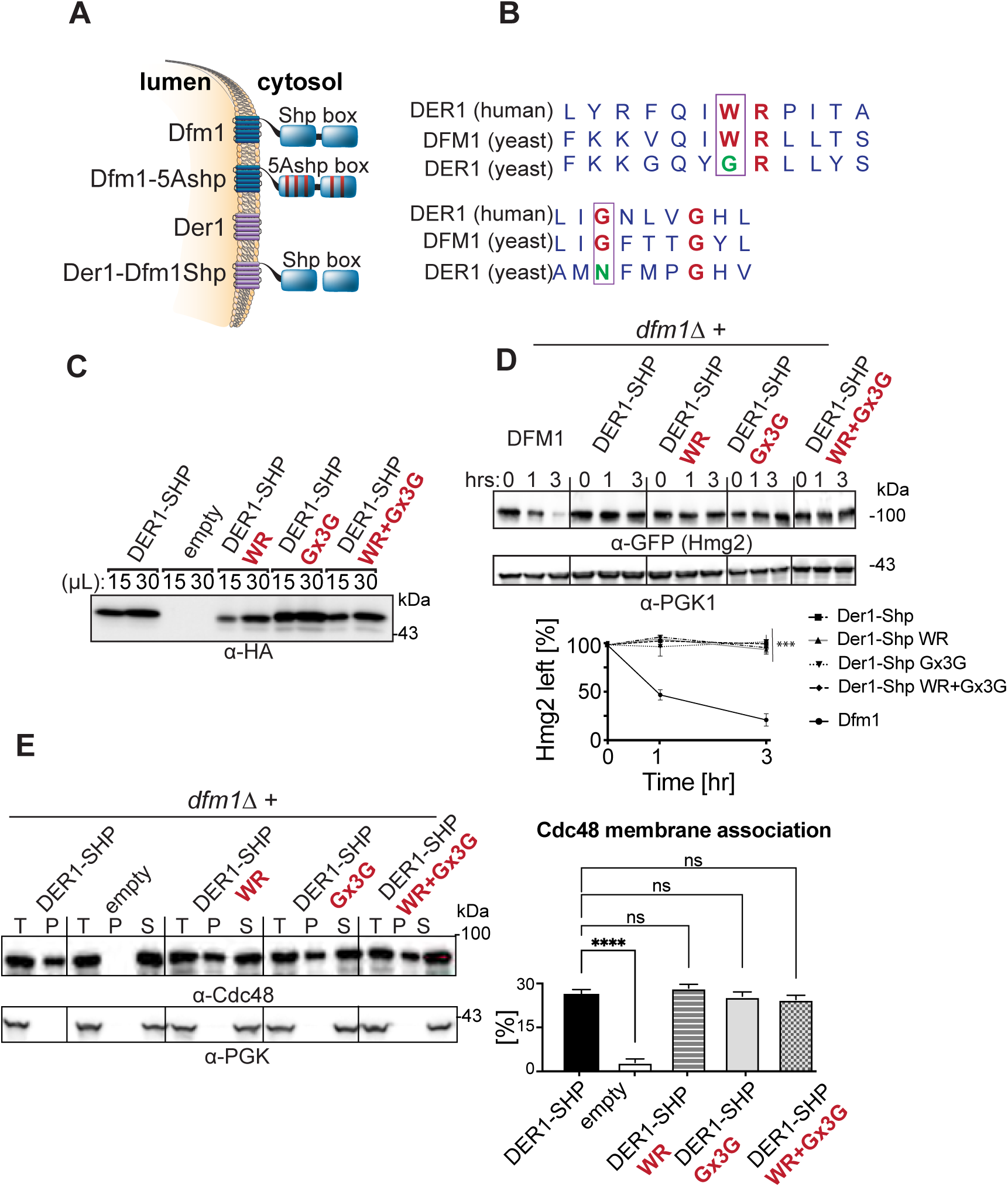
WR and GxxxG motifs are not sufficient for ERAD-M retrotranslocation. (A) Depiction of Dfm1, Der1, and Der1-Shp. Dfm1 and Der1 are ER-localized membrane proteins with six transmembrane domains. Unlike Der1, Dfm1 has an extended cytoplasmic tail containing two SHP boxes. (B) Alignment of WR and GxxxG motif from *H. sapiens* Derlin-1 and *S. cerevisiae* Der1 and Dfm1. (C) Expression levels of Der1-Shp variants are measured by loading increasing amounts of lysates (15 μL and 30 μL) on SDS-PAGE followed by immunoblotting with α-HA. (D). The WR and GxxxG motifs within Der1-Shp are not sufficient for Hmg2 degradation. In the indicated strains, degradation of Hmg2-GFP was measured by CHX-chase assay. Cells were analyzed by SDS-PAGE and immunoblotted for Hmg2-GFP with α-GFP. Band intensities were normalized to PGK1 loading control and quantified by ImageJ. t=0 was taken as 100% and data is represented as mean ± SEM from at least three experiments, ***p < 0.001, Repeated Measures ANOVA. (E) The WR and GxxxG motifs within Der1-Shp does not disrupt Cdc48 recruitment. Total cell lysate (T) from the indicated strains were separated into soluble cytosolic fraction (S) and pellet microsomal fraction (P) upon centrifugation at 14,000 x g. Each fraction was analyzed by SDS-PAGE and immunoblotted for Cdc48 with α-Cdc48 and Pgk1 with α-Pgk1. The graph shows the quantification of Cdc48 in the pellet fractions of the respective cells as measured from ImageJ. Data is represented as percentage of Cdc48 that is bound to pellet fraction and is shown as mean ± SEM from three independent experiments, **** p< 0.0001, One-way ANOVA.

### Dfm1 L1 and TM2 mutants are unable to retrotranslocate ER membrane substrates

To identify additional residues required for Dfm1’s retrotranslocation function, we performed a sequence analysis screen in which random mutagenesis was performed on Dfm1’s transmembrane segment. We excluded mutagenic alteration of the cytoplasmic SHP tail region to prevent false positives due to the disruption of Dfm1’s Cdc48 recruitment function. Dfm1 was mutagenized by using GeneMorph II random mutagenesis kit. Mutagenized Dfm1 was introduced to *dfm1Δ hrd1Δ* null yeast cells containing an optical, self-ubiquitinating substrate, SUS-GFP, a substrate used in our previous screen for discovery and study of Dfm1 retrotranslocation (Fig. 2A,B)(Neal et al., 2018). Notably, because Hrd1 has been shown to be required for restoring retrotranslocation function when Dfm1 is absent (Neal et al., 2018, 2020), our screening strain also has Hrd1 missing to prevent suppression of strains during the random mutagenesis screen. The resulting transformants were screened for high colony fluorescence due to buildup of SUS-GFP, indicating Dfm1 loss of function and inability to retrotranslocate SUS-GFP. Plasmids were extracted from yeast transformants exhibiting high fluorescence and sequenced to discern the causative Dfm1 mutations. Dfm1 mutants containing no premature stop codons and harboring a single mutation were subjected to further analysis. Less interesting possibilities for the loss of Dfm1 function include low expression, incorrect cellular localization, or inability to recruit Cdc48. Accordingly, successful Dfm1 candidate mutants were analyzed for their stability, and if they were robustly expressed, they were then subjected to cellular fractionation and Cdc48 binding assays to exclude mutants that would not elucidate the mechanism of Dfm1. Random mutagenesis was performed near saturation since we were able to recover the same mutants within Dfm1’s TM region. Specifically, we recovered and identified mutants showing robust expression, correct ER localization and no disruption in Cdc48 recruitment activity (Fig. S1B, C, D). Dfm1 was also found to be associated with the E3 ligase HRD complex (Stolz et al., 2010). Through co-immunoprecipitation, we validated that all five mutants did not disrupt their association with major components of the HRD complex, which includes E3 ligase Hrd1 and its partner protein, Hrd3 (Fig. S1E). Next, we generated a structural model of Dfm1 using homology modeling on Der1; an isoform of Dfm1 whose structure has been solved (Wu et al., 2020). Based on this homology model, Dfm1 mutants were enriched in both L1(F58S, L64V and K67E) and TM2 (Q101R and F107S respectively) (Fig. 2C,D & Fig. S1A, *blue asterisks*). Interestingly, Dfm1’s L64 and K67 are conserved across the rhomboid family whereas F107 is conserved specifically among derlins (Fig. S1A).

**Figure 2.**
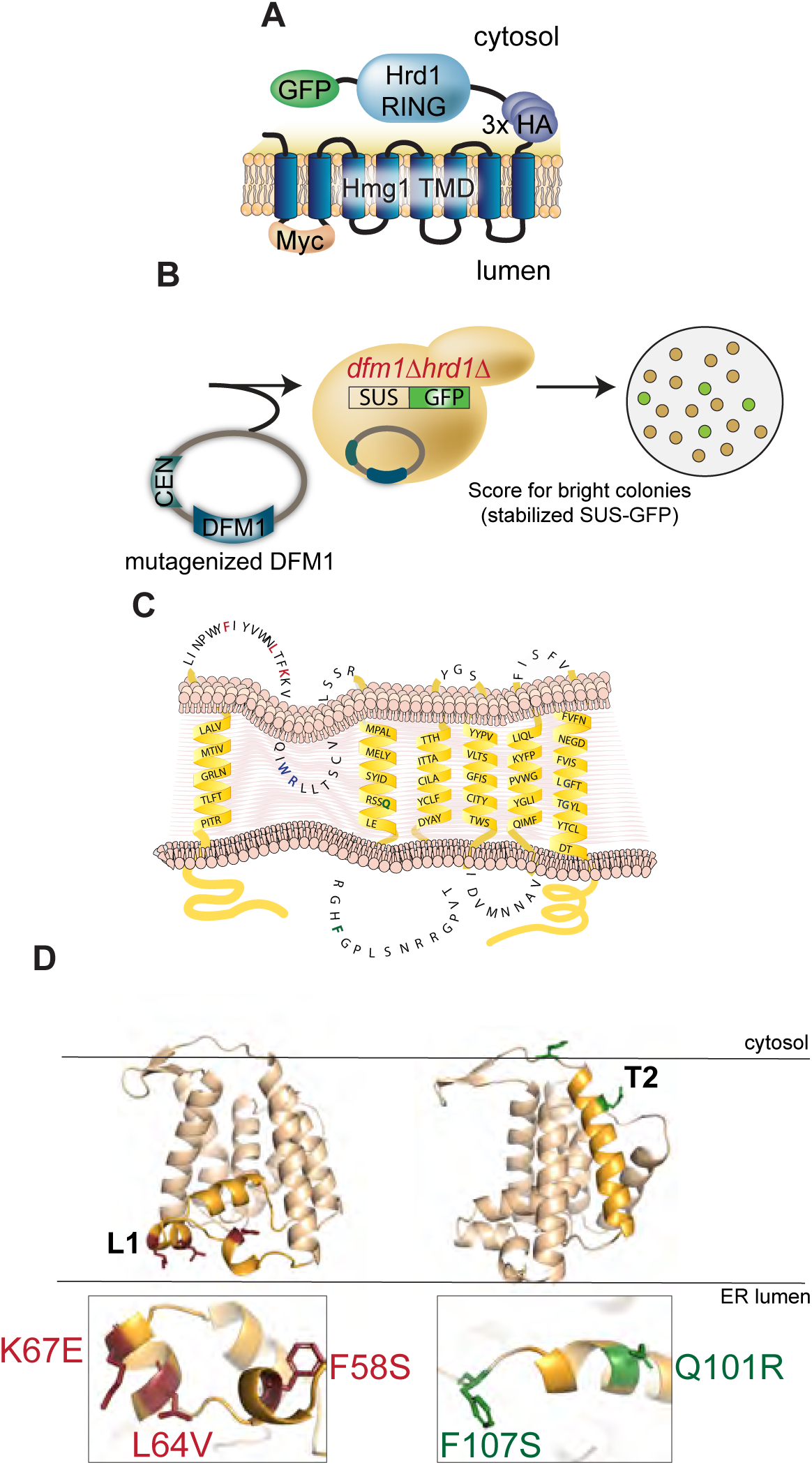
Dfm1 is intolerable to mutations in Loop 1 and Transmembrane domain 2. (A) Depiction of fusion protein, SUS-GFP. The transmembrane Hmg1 domain has a lumenal Myc epitope and the cytosolic domain has three HA epitopes followed by the HRD RING domain fused with the GFP epitope. (B) Full-length DFM1 (except for region encoding SHP box) was subjected to random mutagenesis. Mutagenized DFM1 was transformed into *dfm1Δ hrd1Δ* cells expressing SUS-GFP and scored for stabilization of SUS-GFP or high colony fluorescence by visualization. (C) Depiction of Dfm1 mutants (indicated in red for L1 and green for TM 2) that were selected from the random mutagenesis screen and validated for expression and ability to localize to ER and recruit Cdc48. (D) Homology model of Dfm1. Positions of Loop 1 and TM2 mutants are indicated in red and green respectively.

### L1 and TM2 mutants affect retrotranslocation and ERAD of ER membranes substrates

We examined the extent to which the Dfm1 mutants affected ERAD of various substrates. Test substrates included all Dfm1-dependent membrane substrates characterized in our previous studies: integral membrane HRD pathway substrates Hmg2 and Pdr5* and integral membrane DOA pathway substrate Ste6*. The effect of Dfm1 mutants was directly tested with cycloheximide (CHX)-chase assay on Hmg2, Pdr5* and Ste6* (Fig. 3A-C). The normally degraded substrates, Hmg2 and Ste6*, were completely stabilized by each Dfm1 mutant. In contrast, we observed partial degradation of Pdr5* with each Dfm1 mutant. This is most likely due to the rapid suppressive nature of Dfm1 mutants, which we previously observed to be triggered by overexpression of several ERAD-M substrates (Neal et al., 2018, 2020). Nevertheless, the Pdr5* degradation rate by Dfm1 mutants was slower compared to Pdr5*degradation by wildtype Dfm1. Overall, Dfm1 mutants isolated from the genetic screen affect degradation of ER membrane substrates.

**Figure 3.**
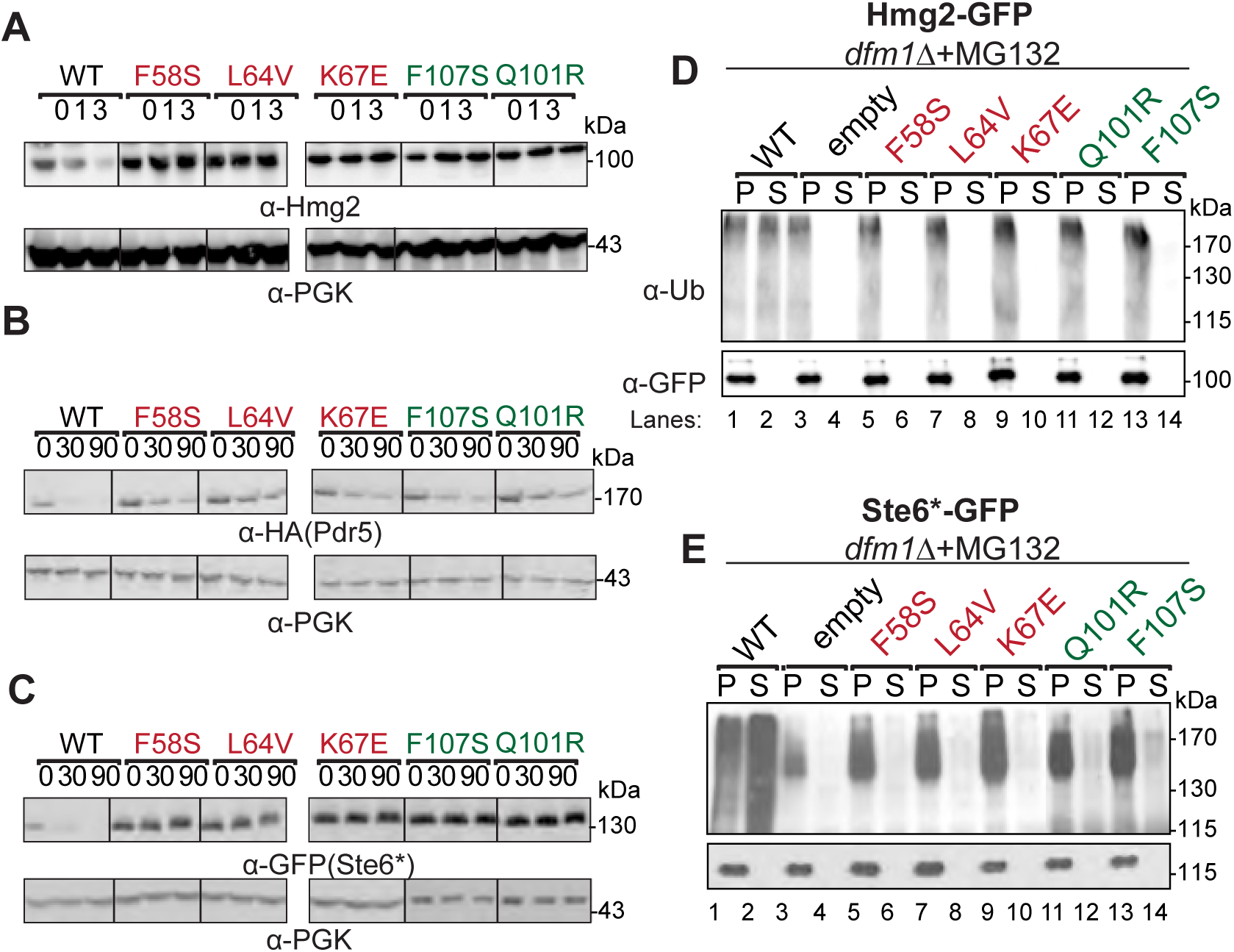
Dfm1 mutants are defective in ERAD-M degradation and retrotranslocation. (A) *dfm1*Δ strains expressing the indicated Dfm1 mutants were grown to log-phase and degradation was measured by CHX. After CHX addition, cells were lysed at the indicated times and analyzed by SDS-PAGE and immunoblotted for Hmg2-GFP with α-GFP. (B) Same as (A) expect degradation of Pdr5*-HA was measured in indicated strains. (C) Same as (A) expect degradation of Ste6*-GFP was measured in indicated strains. (D) Dfm1 mutants are required for retrotranslocation of Hmg2-GFP. Indicated strains were grown to log-phase and treated with MG132 prior to lysis. Crude lysate was prepared from each strain and ultracentrifuged to discern ubiquitinated Hmg2-GFP that either has been retrotranslocated into the soluble fraction (S) or remained in the membrane (P). Following fractionation, Hmg2-GFP was immunoprecipitated from both fractions, resolved on 8% SDS-PAGE and immunoblotted with α-GFP and α-Ubi. (E) Same as (D) except in vivo retrotranslocation assay was performed on Ste6*-GFP.

We confirmed that Hmg2 and Ste6* retrotranslocation was strongly blocked by all Dfm1 mutants in the *in vivo* retrotranslocation assay (Fig. 3D, E). Cells with wild-type functional Dfm1 showed normal Hmg2 and Ste6* retrotranslocation, as indicated by buildup of ubiquitinated Hmg2 and Ste6* in the supernatant (S) fraction, due to inhibition of proteasome function by MG132. In contrast, identical expression of each Dfm1 mutant resulted in buildup of ubiquitinated Hmg2 and Ste6* in the microsomal pellet (P) fraction. This inhibition in retrotranslocation is comparable to controls strains with Dfm1 absent. Thus, by all conditions examined, Dfm1 L1and TM2 mutants are dysfunctional in ERAD and retrotranslocation.

### Dfm1 binds specifically to membrane substrates and not lumenal substrates

Our previous studies showed that Dfm1 is selective for retrotranslocating membrane substrates and not lumenal substrates such as CPY*, KHN and KWW(Neal et al., 2018). This implied that Dfm1 selectively binds membrane substrates and not lumenal substrates. To test this, we directly examined Dfm1 interaction with various classes of ERAD substrates. We analyzed Dfm1 interactions with its well characterized integral membrane substrates: Hmg2, Pdr5* and Ste6*. Hmg2-GFP, Pdr5*-Myc and Ste6*-GFP were immunoprecipitated with GFP or Myc Trap antibodies from lysates of various strains co-expressing Dfm1-HA, followed by SDS-PAGE and immunoblotting for Dfm1 with *α*-HA, Hmg2, and Ste6* with *α*-GFP, and Pdr5* with *α*-Myc (Fig. 4A & S2A,B). In all cases, binding of membrane substrates to Dfm1 was clear. As a control, we tested the very similar, but uninvolved, Der1 homologue for binding to Hmg2-GFP, Pdr5*-Myc and Ste6*-GFP, and found no association (Fig. 4A & S2A,B). We similarly tested Dfm1 interaction with a classical lumenal ERAD-L substrate CPY*. CPY*-GFP was immunoprecipitated with GFP Trap antibodies from lysates co-expressing Dfm1-HA, followed by immunoblotting for Dfm1 with *α*-HA and CPY* with *α*-GFP. We did not observe association of Dfm1 with CPY* whereas association was seen with Der1, its canonical substrate (Fig S2C). These results suggest Dfm1 interacts specifically with ER membrane substrates, but not lumenal substrates.

**Figure 4.**
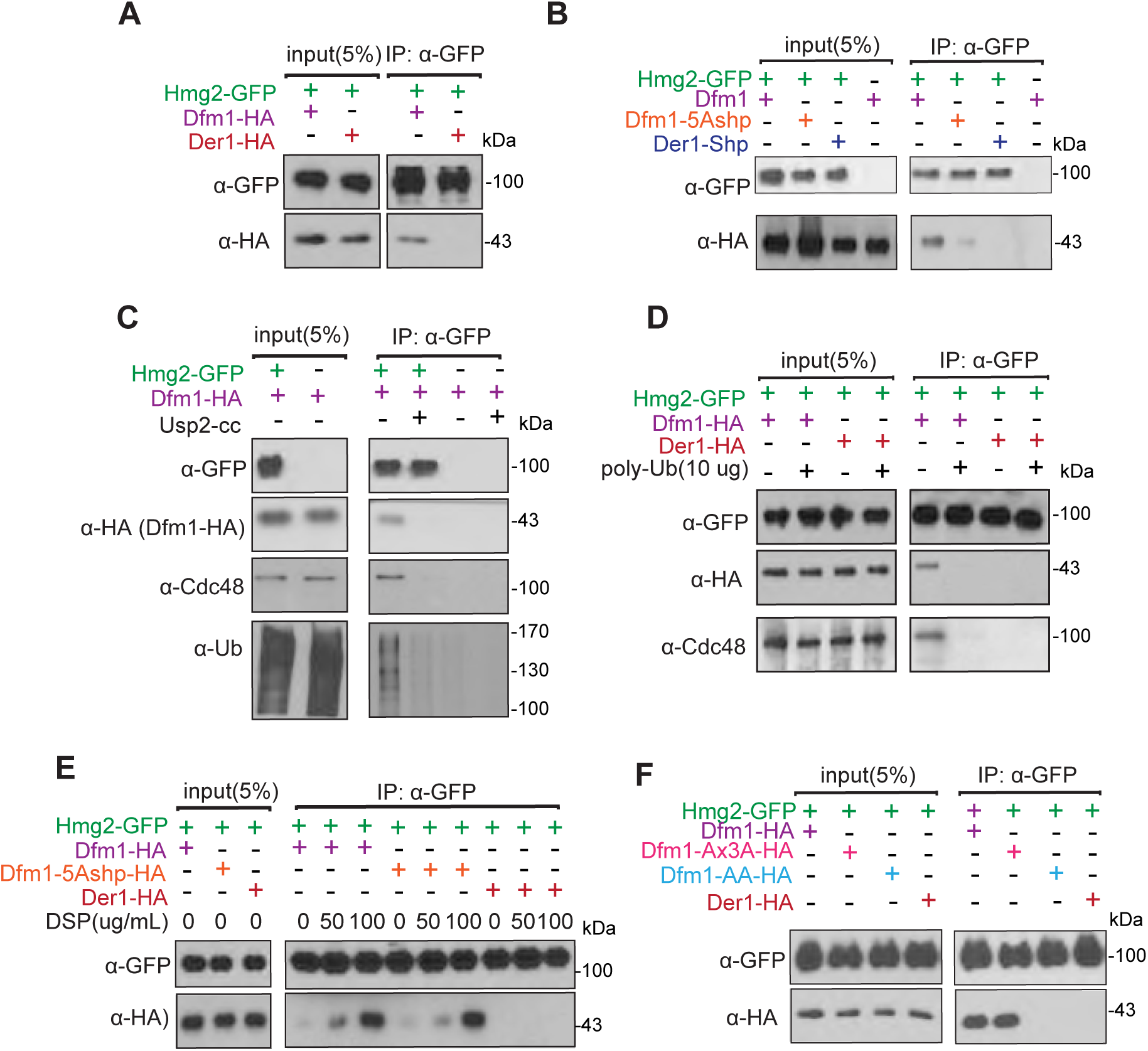
Dfm1’s WR motif and SHP box is required for interaction with membrane substrates. (A) Hmg2-GFP and Dfm1-HA binding was analyzed by co-IP. The IP was analyzed for presence of Hmg2-GFP. As a control for specificity, cells expressing Der1-HA was used. (B) Same as (B) co-IP to analyze binding of Hmg2-GFP to Dfm1 variants: Dfm1-5Ashp and Der1-Shp. As a control for specificity, Der1 was analyzed for binding to Hmg2. (C) Dfm1-Cdc48 complex interacts with polyubiquitin chain of Hmg2. Microsomes isolated from indicated strains were treated with Usp2Core and Hmg2-GFP was immunoprecipitated, resolved on 8% SDS-PAGE and immunoblotted for ubiquitin with α-Ub, Hmg2-GFP with α-GFP, Cdc48 with α-Cdc48 and Dfm1 with α-HA. (D) Addition of Lys48-linked polyubiquitin chains disrupt binding of Hmg2 to Dfm1-Cdc48. Hmg2-GFP, Dfm1-HA and Cdc48 binding was analyzed by co-IP in the presence of increasing amount of Lys48-linked polyubiquitin chains (2 μg, 5 μg and 10 μg). As a negative control, strains not expressing Hmg2-GFP were used. (E) Crosslinking analysis of Hmg2-GFP and Dfm1-5Ashp. Microsomes were harvested from DSP-treated strains and subjected to immunoprecipitation of Hmg2-GFP with GFP Trap, followed by immunoblotting for Dfm1-5Ashp with anti-HA and Hmg2 with anti-GFP. (F) Same as (A), except binding to Hmg2-GFP was analyzed with Dfm1 variants: Dfm1-AA and Dfm1-Ax3A.

### Shp tail is necessary, but not sufficient for substrate binding

We have previously shown that alteration of the 5 signature residues of the Dfm1 SHP box to alanine (Dfm1-5Ashp) removed its ability to recruit Cdc48 (Fig. 1A). (Neal et al., 2018; Sato and Hampton, 2006). Conversely, we have shown that addition of the Dfm1 SHP motif to the normally SHP-less Der1 made this chimera able to promote Cdc48 recruitment comparable to Dfm1, but was not sufficient for supporting retrotranslocation (Fig. 1A)(Neal et al., 2018). We tested these SHP variants for their ability to bind substrates Hmg2, Pdr5* and Ste6*. Notably, the Dfm1-5Ashp mutant that failed to recruit Cdc48 displayed marked disruption in substrate binding with only a small fraction of Dfm1-5Ashp (<5%) bound to ERAD-M Hmg2-GFP, Pdr5*-GFP and ERAD-C Ste6*-GFP compared to wildtype Dfm1 suggesting requirement of SHP tail for substrate association (Fig. 4B & S2A,B). Furthermore, the Der1-SHP chimera that recruited Cdc48, could not support binding to Hmg2, Pdr5* and Ste6*. Thus, the SHP tail, and presumably Cdc48 recruitment to the ER membrane, is necessary, but not sufficient, for substrate association.

### Dfm1 recruits Cdc48 to bind to polyubiquitin chains conjugated to membrane substrates

The requirement for Dfm1’s SHP motif suggests Cdc48 is also involved with binding membrane substrates. Previously, we demonstrated that Dfm1 mediates Cdc48 recruitment to the ER surface(Neal et al., 2018). Moreover, we established that Cdc48 functions as a “retrochaperone” by directly binding to the polyubiquitin chain of membrane substrates to maintain solubility and prevent aggregation of substrates retrotranslocated into the aqueous cytosol(Neal et al., 2017). Based on these observations, we hypothesize that Dfm1 recruits Cdc48 and concomitantly, Cdc48 attaches to the polyubiquitin chains of membrane substrates targeted for retrotranslocation and degradation. We tested whether ubiquitin removal from membrane substrates cause disassociation of substrates from the Dfm1-Cdc48 complex. We used the Usp2Core ubiquitin protease to remove the multiubiquitin chain from ubiquitinated Hmg2-GFP embedded in microsomal membranes. Removal of the ubiquitin chain from Hmg2-GFP caused loss of both the covalently bound ubiquitin and the associated Dfm1-Cdc48 complex (Fig.4C). Because the Dfm1-Cdc48 complex binds polyubiquitin chains, we next tested the effect of adding commercially available free polyubiquitin chains on the association of Dfm1-Cdc48 complex with Hmg2-GFP. Direct addition of increasing amounts of Lys-48-linked polyubiquitin to microsome fractions completely blocked the association of Dfm1-Cdc48 with Hmg2-GFP as assessed by GFP-Trap coprecipitation (Fig. 4D). We interpreted this to mean that ubiquitinated Hmg2 is removed by direct competition of free polyubiquitin chains.

### WR motif is required for membrane substrate binding

The above studies show that Dfm1’s SHP tail—through Cdc48 binding—is required for direct interaction with polyubiquitinated membrane substrates. We observed no association of membrane substrate with chimera Der1-SHP with intact Cdc48 recruitment function suggesting Cdc48 is not sufficient for substrate interaction (Fig. 4B). Furthermore, a small fraction of Dfm1-5Ashp without intact Cdc48 recruitment function was found to be associated to Hmg2-GFP, suggesting a transient interaction between Dfm1-5Ashp and Hmg2 (Fig. 4B). Indeed, we observed stable association of Hmg2-GFP with Dfm1-5Ashp with crosslinking, confirming that the interaction is transient and suggests substrate binding is mediated by additional information within the Dfm1 transmembrane region and is independent of ubiquitin binding (Fig. 4E). Accordingly, we analyzed whether additional residues within Dfm1’s transmembrane domains are required for membrane substrate interaction. Previously, we have shown that Dfm1’s conserved rhomboid motifs WR and GxxxG are required for retrotranslocation(Neal et al., 2018). We utilized two mutants from our previous studies, Dfm1-AA and the Dfm1-AxxxA in which the conserved residues in the WR or the GxxxG motif were mutated to alanine. Both Dfm1-WR/AA and Dfm1-GxxxG/AxxxA were employed in the substrate binding assay in cells expressing Hmg2, Pdr5* or Ste6* (Fig. 4F & S2A,B). As a control for specificity, we tested Der1 binding to all three membrane substrates, which are not its substrate, and found no detectable association. Binding of Dfm1-AxxxA to Hmg2, Pdr5* or Ste6* was clearly detectable and to the same extent found for binding to wildtype Dfm1. In contrast, there was no association of Dfm1-WR/AA to membrane substrates (Fig. 4F & S2A,B). These results indicated that the WR, and not GxxxG, motif is required for binding to ER membrane substrates tested.

### Dfm1 L1 retrotranslocation-deficient mutants are unable to bind ER membrane substrates

Results above indicate the presence of additional residues along Dfm1’s transmembrane region that is involved with substrate binding. From our sequence analysis, we have identified Dfm1 L1 and L2/TM2 mutants that affect ERAD and retrotranslocation of a variety of ER membrane substrates. We next examined whether the L1 and TM2 mutants affect membrane substrate detection. We employed the substrate binding assay to analyze association of Dfm1 mutants with various membrane substrates: Hmg2-GFP, Pdr5*-Myc and Ste6*-GFP (Fig. 5A-C). Each substrate was immunoprecipitated with GFP or Myc Trap followed by SDS-PAGE and immunoblotted for Dfm1 with *α*-HA, Hmg2, and Ste6* with *α*-GFP, and Pdr5* with *α*-Myc. In all cases, membrane substrates were still associated with Dfm1 TM2 mutants to an extent similar to binding to wildtype Dfm1. In contrast, there was no detectable association of membrane substrates with Dfm1 L1 mutants, implying that\ all three L1 residues are required for membrane substrate binding.

**Figure 5.**
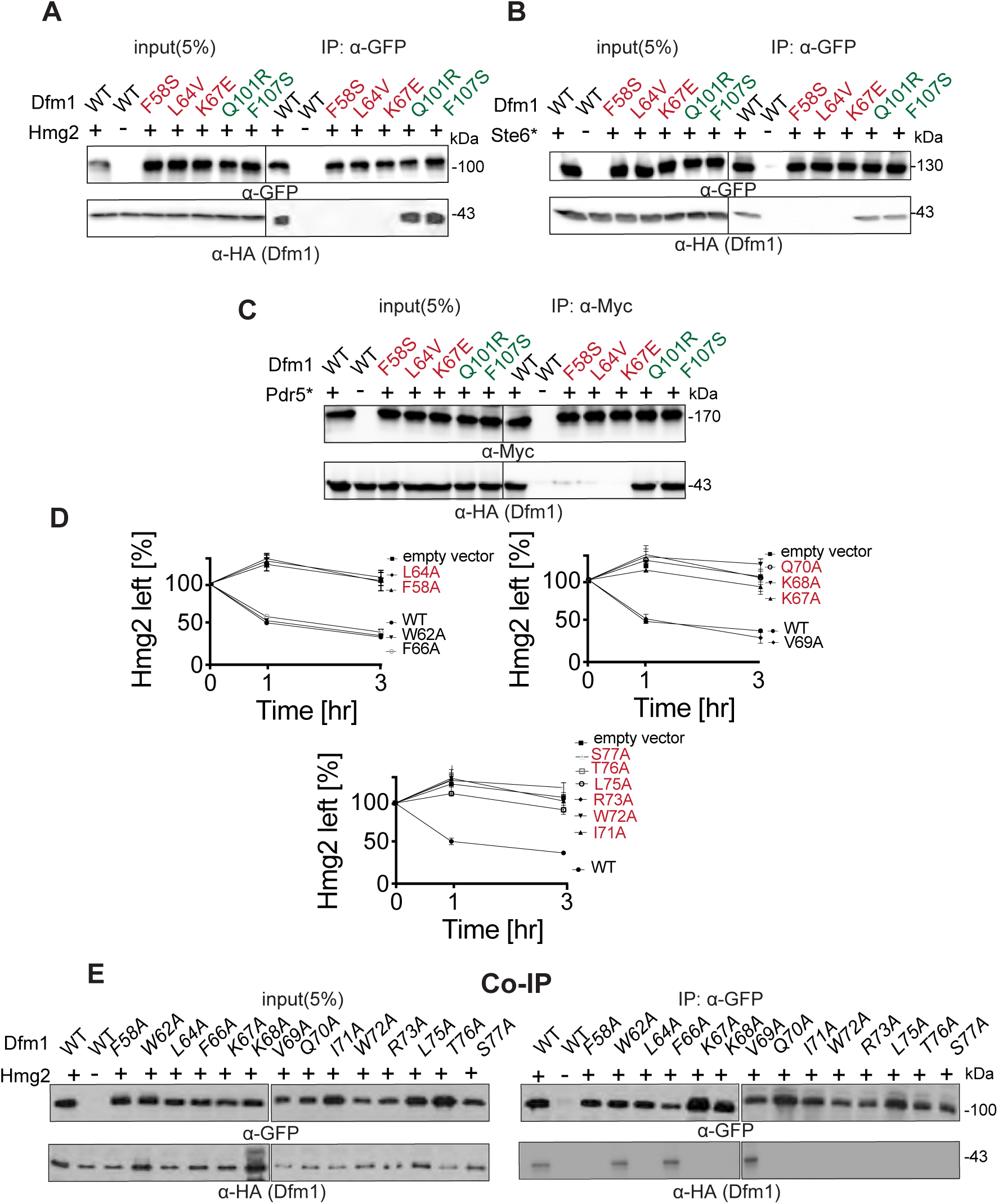
Dfm1 L1 residues are required for binding to integral membrane substrates. (A) Hmg2-GFP and binding to retrotranslocation-deficient Dfm1 mutants was analyzed by co-IP. The IP was analyzed for presence of Hmg2-GFP. As a negative control, cells not expressing Hmg2-GFP were used. (B) Same as (A) except Ste6* and binding to retrotranslocation-deficient Dfm1 mutants was analyzed by co-IP. (C) Same as (A) except Pdr5* binding to retrotranslocation-deficient Dfm1 mutants was analyzed by co-IP. (D) Flow cytometry shows a subset of L1 Dfm1 mutants generated by Ala mutant scanning strongly blocks Hmg2 ERAD. The indicated strains were grown to log phase and were subjected to CHX-chase. Hmg2-GFP levels were analyzed at the indicated times using flow cytometry. Histograms of 10,000 cells are shown, with the number of cells *versus* GFP fluorescence. Data from each time point is represented as mean ± SEM from at least three experiments. (E) Same as (A) except Hmg2-GFP binding to L1 Dfm1 mutants generated by Ala mutant scanning were analyzed by Co-IP.

This result suggests a critical role for L1 in membrane substrate binding, which is in agreement with previous reports that L1 region in GlpG, a bacterial rhomboid, plays a critical role in substrate engagement (Zoll et al., 2014). Although sequence similarities across the rhomboid superfamily are low, there are highly conserved residues that cluster in the L1 region (Fig. S1A). Based on GlpG’s crystal structure, L1 is embedded in the lipid bilayer, a feature that is uncommon amongst membrane proteins (Lemieux et al., 2007; Wang et al., 2006). The uniqueness of this motif prompted us to investigate L1 in more detail. Accordingly, we performed mutagenesis on the L1 region in which each residue was mutated to Alanine. To rule out the possibility that mutants resulted in no expression or mislocalization of Dfm1, we performed western blotting on Dfm1 levels (Fig.S3A). Notably, mutations within a hydrophobic patch of L1 (P55A, W56A, Y57A, I59A, Y60A, and V61A) resulted in no expression of Dfm1(Fig. S3A,C,D). As expected, the only mutant within this cluster, which was expressed at similar levels to wildtype, was F58A; the original mutant recovered from our mutagenesis screen. Based on lack of expression, it appears this hydrophobic patch plays a critical in the structural stability and/or membrane insertion for Dfm1(Fig.SD). Furthermore, Dfm1 mutants N63A, T65A, and L74A resulted in no expression and hence these mutants were also excluded for further functional analyses. All other mutants showing robust Dfm1 expression and exhibiting correct localization (Fig. S3B) were analyzed for their effect on the steady-state levels of the self-ubiquitinating substrate, SUS-GFP and ERAD of Hmg2-GFP (Fig. 5D & S3C). As measured by flow cytometry, F58A, L64A, K67A, K68A, Q70A, I71A, W72A, R73A, L75A resulted in high steady-state levels of SUS-GFP and a strong block in Hmg2-GFP degradation with GFP levels that were comparable to control cells lacking Dfm1. Notably, these same residues resulted in the inability to bind to membrane substrate, Hmg2 (Fig. 5E). In contrast, W62A, F66A, and V69A mutants didn’t disrupt Dfm1-mediated degradation of SUS and Hmg2 and were able to support substrate binding. Overall, mutants disrupting Dfm1’s action mapped to sites that are highly conserved in the L1 region amongst the rhomboid superfamily, further validating a critical role for L1 in substrate binding.

### Dynamic interaction of Dfm1 and the lipid bilayer

Recent work demonstrated Dfm1’s homolog, Der1, forms a half channel with E3 ligase Hrd1 to induce lipid thinning, which facilitates in the retrotranslocation of lumenal ERAD-L substrates (Wu et al., 2020). We hypothesize Dfm1 has retained membrane perturbation properties to aid in the removal of multi-spanning membrane substrates. To examine this, we first built a homology model of Dfm1, using the recently solved structure of its homolog, Der1, as a template structure(Wu et al., 2020). Molecular dynamics (MD) simulations were performed to examine the lipid interactions of Dfm1 embedded in a mixed lipid bilayer representative of the ER membrane. Lipid thickness in distant regions from Dfm1 was approximately 4.0-4.5 nm, which is expected for phospholipid bilayers (Bondar, 2020). In contrast, we observed rearrangement of lipids in the vicinity of Dfm1, near TM1, TM2 and TM5 (Fig. 6A,B). Lipids were perturbed on both the luminal and cytoplasmic side, and lipid thinning was observed with a membrane thickness of approximately 2.0-2.5 nm (Fig. 6A,B,C, *circled in blue*). Furthermore, through the duration of the simulation, local lipid thinning remained in the same **region**, in between TM 2&5 and between TM 1&2 (Video Fig. 1). Local lipid thinning of this magnitude has also been reported to occur in the same region of Dfm1’s paralog, Der1(Wu et al., 2020). Interestingly, the region of membrane thinning (between Dfm1’s TM 2 and 5) is localized in an area that was determined to be the lateral gate for bacterial and yeast rhomboid, GlpG and Der1 respectively. (Fig.6A, indicated with asterisk) (Lemieux et al., 2007; Wang et al., 2006; Wu et al., 2020)

**Figure 6.**
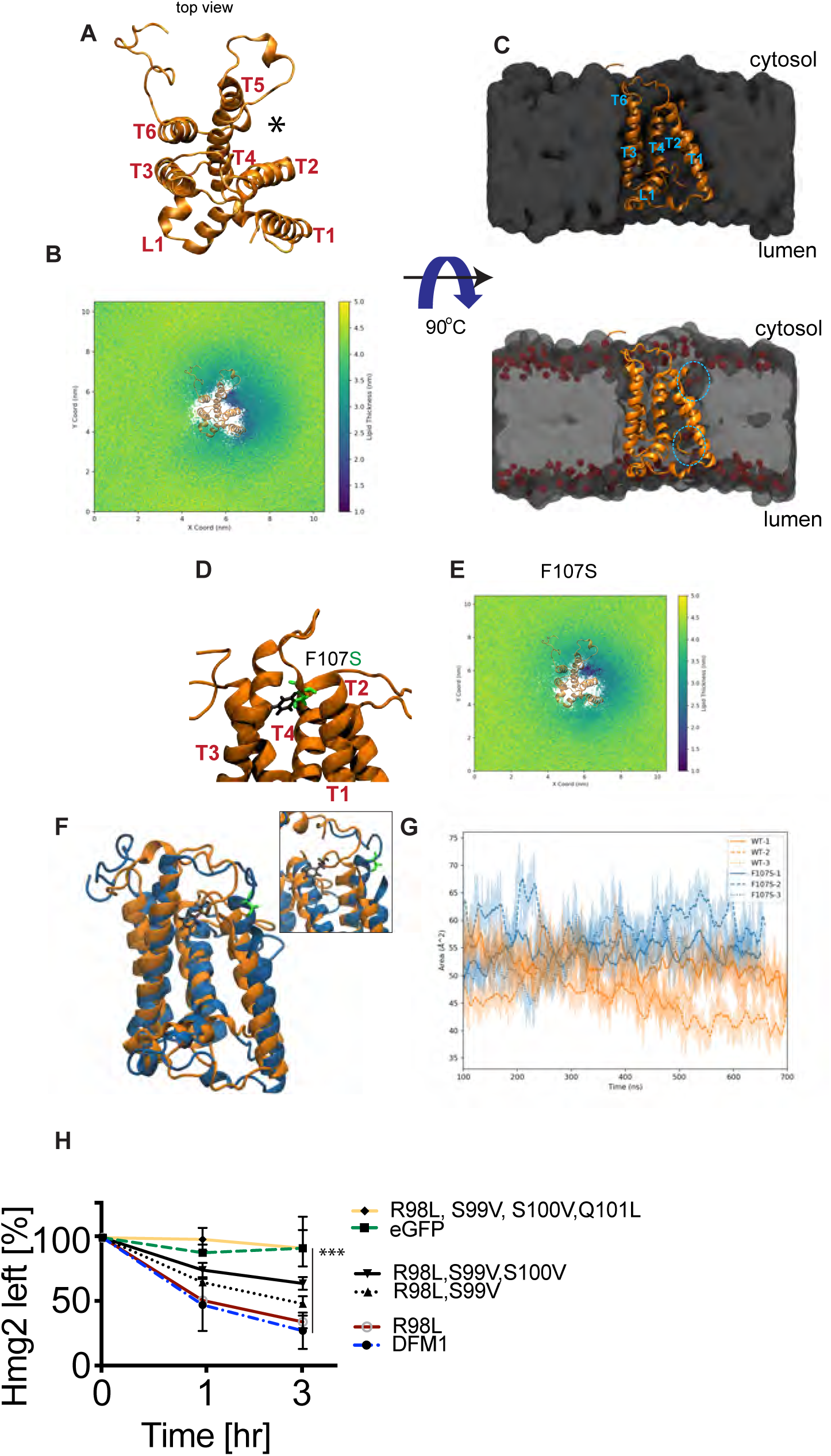
Membrane thinning by Dfm1. (A) Top view of *S. cerevisiae* derlin, Dfm1, homology model shown in gold ribbon with respective transmembrane domains and Loop 1 labeled. (B) Membrane thickness is shown as x and y 2D maps of the positions of the lipid head groups every 1ns of simulation color-coded with the membrane thickness at that timepoint/location. The Dfm1 protein model is overlayed to show the relative locations of membrane thinning. Total thickness, i.e. the distance calculated between the upper and lower surfaces used for the analyses, is shown color-coded according to a 1.0 to 5.0 nm range. (C) Midpoint cross-section of the membrane where Dfm1 is embedded. Dfm1 is shown in gold ribbon, the lipids are shown in a grey volumetric representation, and the phospholipid head group is shown in red. (D) Homology model of Dfm1 with residues F107 shown in black and mutant residue F107S shown in green. (E)As in (B), except membrane thickness was analyzed for Dfm1 mutant F107S. (F) Protein structure clusters with the highest prevalence (∼50% of simulation time) of the WT protein (gold) and F107S protein (blue), highlighting the residue positional difference in F107 (black) and F107S (green). (F) SASA plots for the MD simulations with the shaded regions showing the raw data and the lines showing the 10 ns moving average. (H) *dfm1*Δ strains expressing the indicated Dfm1 mutants were grown to log-phase and degradation was measured by CHX-chase analysis. After CHX addition, Hmg2-GFP levels were measured by flow cytometry cells. Data is represented as mean ± SEM from three experiments, ***p < 0.001, Repeated Measures ANOVA. (I) As in (B), except membrane thickness was analyzed for Dfm1 quad mutant, R98L, S99V, S100V, Q101L.

### Dfm1 TM2 mutants disrupt lipid thinning activity

The observed lipid deformation near TM1, TM2 and TM5 indicates an important role of these transmembrane helices in lipid thinning and retrotranslocation. We have isolated a mutant from our random mutagenesis screen, TM2 residue F107S, which is near the site of lipid perturbation (Fig. 6C,D).This mutation has been validated in our assays for disrupting membrane substrate retrotranslocation, but do not disrupt substrate binding (Fig. 3A-C & 5A-C) To investigate how this mutant may disrupt lipid thinning and retrotranslocation, MD simulations were performed with the F107S mutant in comparison to the wildtype. Simulations with the F107S mutation effected lipid thinning in the vicinity of TM1 and TM2. Wildtype Dfm1 had a more consistent thinning effect around TM1, TM2, and TM5 where the membrane tended to be around 2.0-2.5 nm, while F107S Dfm1 had a comparable thinning effect around TM2 and TM5, but a reduced thinning effect near TM1 at approximately 3.5-4.0 nm. (Fig. 6E). The F107S mutant also significantly altered the structure of Dfm1 and increased the solvent accessible surface area (SASA) of the protein while having negligible effects on the total exposed surface area of the protein (Fig. 6G & S4). Accordingly, F107 appears to play a structural role and mutation of this residue affects Dfm1’s lipid perturbation properties and renders Dfm1 completely dysfunctional in retrotranslocation (Fig. 3E,F & 6E).

Dfm1’s isoform, Der1, contains hydrophilic stretches at TM2 (residues NHLST) that are critical for lipid thinning via their interactions with the phosphate head group of the lipid bilayer (Wu et al., 2020). Dfm1 contains an analogous cluster of hydrophilic residues (RSSQ). Interestingly, Q101R –the retrotranslocation-deficient Dfm1 mutant isolated from random mutagenesis– is within this hydrophilic cluster (RSS**<u>Q</u>**). To test the functional importance of the hydrophilic TM2 residues, we mutated these residues to hydrophobic amino acids in order to increase the hydrophobicity of TM2. The single, double, triple and quadruple mutants were localized correctly in the microsome fraction (Fig. S4D) and each mutant reduced the degradation rate of Hmg2, with strongest stabilization with the quadruple mutant: (R98L, S99V, S100V, Q101L) (Fig. 6H). These findings suggest that Dfm1’s mechanistic action for lipid thinning is analogous to Der1.

### Derlin-1 homologous mutants disrupt ERAD of CFTR

The studies above have identified sequence features of yeast rhomboid, Dfm1, that are important for its retrotranslocation function via membrane substrate detection and lipid thinning. The closest human homolog to Dfm1 is Derlin-1; the most well characterized human derlin to date (Greenblatt et al., 2012; Sun et al., 2006a; Suzuki et al., 2012; You et al., 2017). In fact, yeast Dfm1 has higher sequence similarity to human Derlin-1 in comparison to its yeast counterpart Der1 (Sato and Hampton, 2006b). Like Dfm1, Derlin-1 possesses a Shp tail for recruiting p97/Cdc48 along with the conserved WR and GxxxG motifs, which have been shown to be critical for its retrotranslocation function(Greenblatt et al., 2011). We wanted to perform a similar random mutagenesis screen on full-length human Derlin-1 by leveraging our established screen in *S. cerevisiae*. We first verified whether full-length human DERLIN-1 or its other ERAD-participating paralog, DERLIN-2 gene can complement the *dfm1Δ* phenotype in *S. cerevisiae*. DERLIN-1 or DERLIN-2 yeast-optimized coding sequence was inserted into the yeast expression plasmid and transformed into *dfm1Δ* cells expressing ERAD-M substrate, Hmg2-GFP. Both Derlin-1 and Derlin-2 steady-state levels were detectable by western blotting (Fig. S5A). However, both Derlin-1 and Derlin-2 were not able to degrade Hmg2-GFP implicating both human derlins are not able to functionally complement yeast derlin Dfm1 (Fig. S5B).

We next employed the mammalian system to examine sequence requirements for human derlin function. Interestingly, a subset of Dfm1 residues that were identified from random mutagenesis and Ala mutant scanning (L1: L64, K68, L75 and TM2: F107S) are similarly conserved in its human homolog, Derlin-1 (L1: A45, R49, I56 and TM2: F91, respectively) (Fig. 7A). We determined whether these conserved Derlin-1 residues with similar properties are critical for ERAD. We performed site-directed mutagenesis on these conserved residues. W53A mutant in the WR motif was also generated as a control, since this motif has been shown to be required for human Derlin-1 retrotranslocation function(Greenblatt et al., 2011). All Derlin-1 mutants displayed robust expression and correct localization to the ER (Fig. 7B,C & S5C). A well characterized multi-spanning membrane substrate for Derlin-1 is the clinically important disease-causing mutant cystic fibrosis transmembrane conductance regulator (CFTR)ΔF508(Sun et al., 2006b). By using a Derlin-1 knockout cell line expressing CFTRΔF508, Derlin-1 mutants were directly tested with CHX-chase assays measuring CFTRΔF508 degradation. Remarkably, in all cases, the normally degraded CFTRΔF508 was stabilized by all Derlin-1 mutants with levels similar to GFP only and WR mutant control (Fig. 7F). Subsequently, we discerned whether the Derlin-1 L1 and TM2 mutants affected their binding to CFTRΔF508. Each Derlin-1 mutant was subjected to pull-down experiments with Ni^2+^ agarose beads. A fraction of CFTRΔF508 (∼5%) copurified with wildtype Derlin-1 (Fig. 7D). As a control for specificity, CFTRΔF508 did not bind to resins in GFP only cells. CFTRΔF508 associated with Derlin-1 TM2 mutant F91A to an extent similar to wildtype Derlin-1. In contrast, there was no detectable association of CFTRΔF508 with Derlin-1 mutants (A45S, R49A, W53A) and decreased association of CFTRΔF508 (∼1%) with Derlin-1 mutant I56A, implying that Derlin-1 L1 residues also contribute to membrane substrate binding of CFTRΔF508. Overall, these results indicate yeast Dfm1’s retrotranslocation-specific residues are conserved and also critical for substrate engagement and lipid thinning function of its human homolog, Derlin-1.

**Figure 7.**
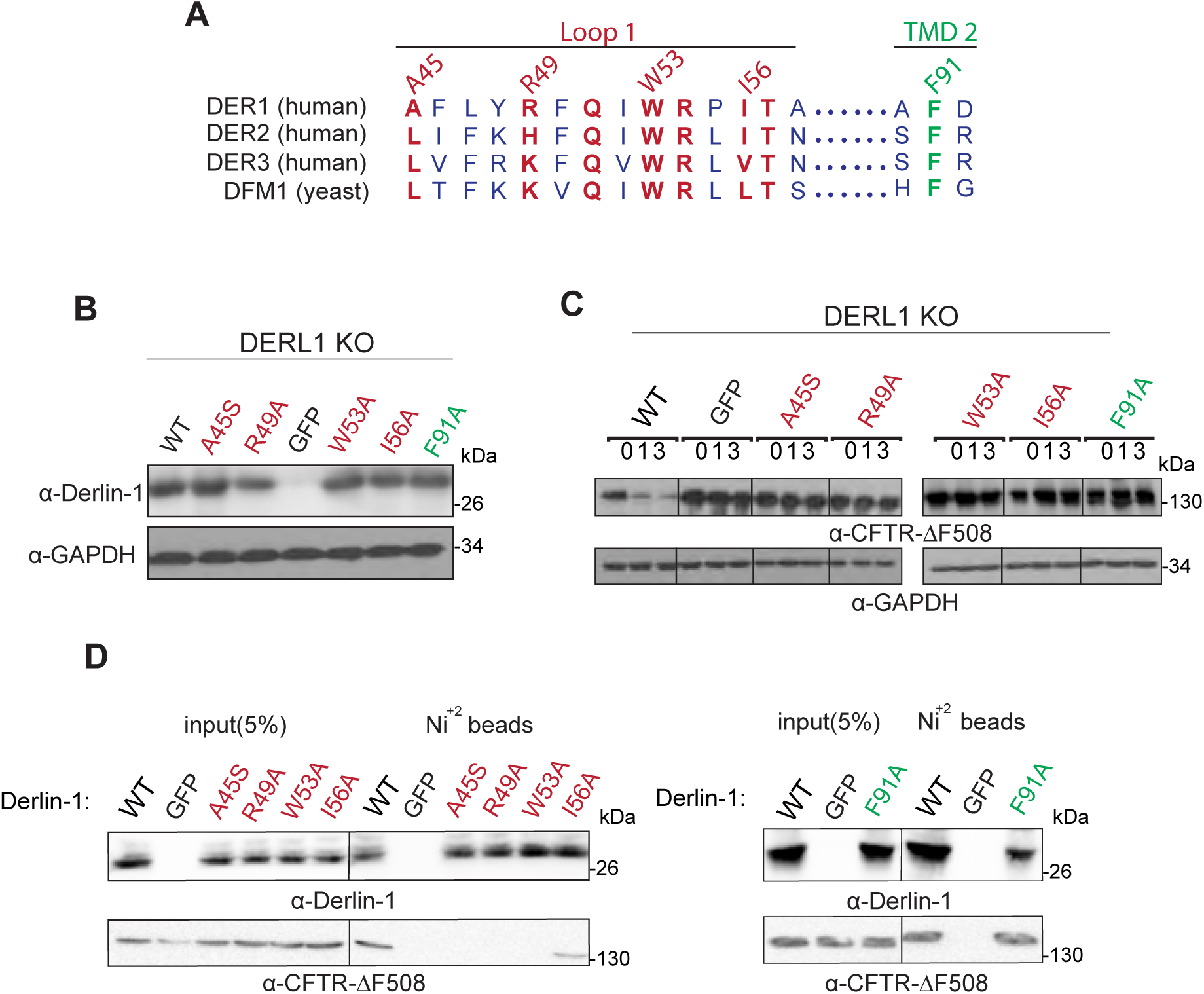
Conservation of human derlin, Derlin-1, ERAD function. (A) Alignment of *H. sapiens* Derlin-1,2,3 and *S. cerevisiae* Dfm1. Similarly or identically conserved residues in loop1 and transmembrane domain 2 are highlighted in red and green respectively. (B) DERLIN-1 KO HEK293T cells were transfected with DERLIN-1 as described under “STARS Methods.” 50 μg of lysate was subjected to immunoblotting for Derlin-1 with α-Derlin-1 and GAPDH with α-GAPDH. (C) Cycloheximide-chase were performed in DERLIN-1 KO HEK293T cells co-transfected with DERLIN-1 and *Δ*F508-CFTR. 48 hours after co-transfections, cells were treated with 100 µg/mL and harvested at the indicated chase times for immunoblotting of CFTR. (D). HEK293T lysates were incubated with Ni^+2^ beads and the bound proteins were eluted with SDS-PAGE sample buffer, resolved by SDS-PAGE and analyzed by immunoblotting with specific antibodies against Derlin-1 and CFTR.

## Discussion

Our studies shed valuable insights on the retrotranslocation function of rhomboid pseudoproteases. How catalytically inactive rhomboid-like proteins are able to retrotranslocate ERAD membrane substrates is an intriguing question in the field. In this study, we unveiled the mechanistic features of derlins, a subclass of rhomboid-like proteins that are widely represented in ERAD. Specifically, we discovered L1 and TM2 residues are critical for yeast derlin Dfm1’s actions. Closer analysis of these retrotranslocation-deficient mutants reveal L1 mutants are defective in detecting membrane substrates, suggesting the L1 regions are required for substrate binding. Our studies also provide the first evidence that rhomboid pseudoproteases, such as yeast derlin Dfm1, have retained membrane-perturbing properties for channeling or moving multi-spanning integral membrane substrates through the lipid bilayer during retrotranslocation. This was confirmed by molecular dynamic simulations on Dfm1’s homology model coupled with genetics and our *in vivo* retrotranslocation assay, demonstrating that Dfm1 possesses lipid deformation properties, causing a TM2 retrotranslocation-deficient mutant to specifically disrupt lipid thinning. Overall, this study establishes that yeast derlin Dfm1 utilizes the unique properties of the rhomboid superfamily for carrying out the widely conserved and critical process of membrane protein retrotranslocation.

Our co-IP experiments with Dfm1 mutants and chimeras indicate substrate detection requires Dfm1’s SHP tail through Cdc48 recruitment. Notably, Cdc48, which is recruited by Dfm1, directly binds to the polyubiquitin chain of substrates. This was evident by treatment with a deubiquitinase in which cleavage of the polyubiquitin chain liberated membrane substrate Hmg2 from the Dfm1-Cdc48 complex. This was further corroborated through the ability of excess polyubiquitin chains to compete with ubiquitinated substrate, Hmg2, for binding to Dfm1-Cdc48. This finding is consistent with previous works in which Cdc48 along with its cofactors Npl4 and Ufd1 binds directly to ubiquitinated membrane substrates that are destined for retrotranslocation and proteasomal degradation (Bodnar and Rapoport, 2017; Neal et al., 2017). Finally, additional studies demonstrate that several members of the rhomboid superfamily recruit Cdc48/p97 to associate directly with the polyubiquitin chains attached to substrates. For example, human rhomboid protease Rhbdl4 and *S. pombe* rhomboid pseudoprotease, Rbd2, recruit Cdc48/p97 through their VBM and SHP motifs, respectively, for direct interaction of polyubiquitin chains on substrates(Fleig et al., 2012; Hwang et al., 2016). Thus, binding to polyubiquitin chains of substrates appears to be a feature utilized by a subset of the rhomboid superfamily.

It also appears that the Dfm1 L1 region mediates the recognition of integral membrane substrates, as demonstrated by our discovery of L1 retrotranslocation-deficient mutants highly selective for ERAD membrane substrates. This is similar to its bacterial homolog, GlpG, which also possesses a widely conserved L1 that protrudes into the lipid bilayer to facilitate in lipid thinning and substrate binding (Ha et al., 2013; Lemieux et al., 2007; Zoll et al., 2014). An important question that arises from our observations is: how does Loop 1 gain substrate access? One possibility is that Dfm1’s L1 region is poised to attract membrane substrates possessing unstable and/or positively charged transmembrane helices. For example, a previous study demonstrated membrane substrates with helix breaking residues and limited hydrophobicity move readily into the hydrophilic interior cavity of GlpG (Moin and Urban, 2012). Additionally, the mammalian rhomboid protease Rhbdl4 has been shown to have a preference for membrane substrates with positively charged transmembrane helices (Fleig et al., 2012). Indeed, MD simulation of Dfm1 with H_2_0 molecules demonstrated that the interior of Dfm1 is hydrophilic, supporting its role in attracting membrane substrates with positively charged transmembrane helices (Fig. S4A). Interestingly, a similar simulation with Der1 demonstrated one side of the protein was hydrophilic, supporting its role in functioning as a “half channel” for transporting soluble membrane substrates (Fig. S4B). Alternatively, L1 can aid in the diffusion of Dfm1 through the lipid bilayer to survey for membrane substrates destined for retrotranslocation (Kreutzberger et al., 2019). This alternative mode of substrate detection is supported from previous work suggesting a subset of the rhomboid family diffuses rapidly through the lipid bilayer in order to patrol through the membrane to allow for substrate detection. Overall, based on requirement of Dfm1 L1 and Dfm1 recruitment of Cdc48 for substrate association, we suggest a model of at least two coordinated functions of Dfm1 in substrate detection. We propose a model in which Dfm1 L1 region brings the substrate in close proximity to Dfm1; allowing for concomitant binding to the polyubiquitin chain attached to the substrate (Fig.8). Nevertheless, understanding the mechanism of substrate detection and association is of high importance and would be significantly advanced by high-resolution structural characterization of substrate binding.

**Figure 8.**
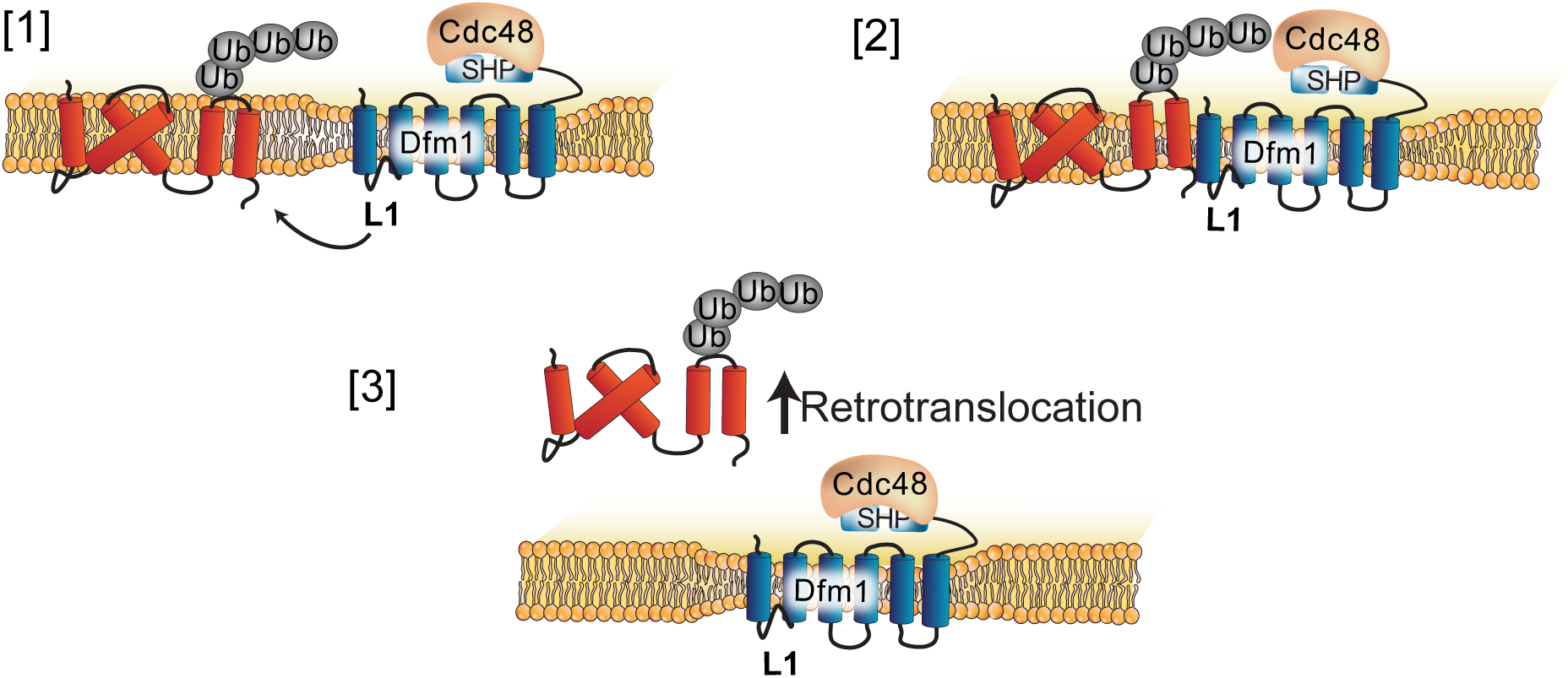
Model of derlin Dfm1-mediated retrotranslocation of integral membrane substrates. [1] Dfm1 targets and binds integral membrane substrates through its L1 region. [2] Once in close proximity, the polyubiquitin chain attached to substrates bind directly to Cdc48 that was recruited by Dfm1. [3] Lipid-thinning function by Dfm1 facilitates in the extraction of integral membrane substrates form the ER.

Cells must have a strategy in place for overcoming the thermodynamic barrier of removing hydrophobic integral membrane proteins from their stable home within the lipid bilayer (Marinko et al., 2019). For example, the magnitude of this energetic barrier has been shown by bacteriorhodopsin, a membrane protein, which exhibits a free energy difference of 230 ± 40 kcal/mol between its native and unfolded state(Müller et al., 2002). Previous studies demonstrate how yeast derlins Dfm1 and Der1 can overcome the energetic barriers associated with retrotranslocation (Neal et al., 2018; Wu et al., 2020). For instance, the unique structure of the rhomboid proteins has been proposed to reduce the thickness of the membrane bilayer, reducing the lipid permeability barrier (Bondar et al., 2009b; Marinko et al., 2019). Recently, it was shown that yeast derlin Der1 associates with Hrd1 to induce lipid thinning and permeabilize the membrane for transport of luminal substrates(Wu et al., 2020). Whether Dfm1 employs the same mechanism for retrotranslocating membrane substrates remained unknown and has recently been attained in our study. Through sequence analysis and molecular dynamics simulation of Dfm1’s homology model, we demonstrated that Dfm1 distorts the surrounding lipid bilayer.

Furthermore, we identified retrotranslocation-deficient mutants that ablate Dfm1’s lipid thinning function. Just like Dfm1’s homolog, Der1, such membrane perturbation activity from Dfm1 would lower the energetic cost of removing membrane substrates (Marinko et al., 2019). In vitro studies by Schmidt et al., has shown that E3 ligase Doa10 is required to dislocate a single-spanning membrane substrate, Ubc6, from the liposome (Schmidt et al., 2020). It is possible, Doa10 also possesses lipid thinning properties and may act in concert with Dfm1 to facilitate in extraction of a subset of multi-spanning membranes with varying degrees of hydrophobicity. Furthermore, Dfm1’s rhomboid predecessors are believed to bind to their substrates within the membrane to partially (and passively) unfold the TM helices of substrates prior to proteolytic cleavage (Moin and Urban, 2012; Wang et al., 2007). Since rhomboid pseudoproteases, such as Dfm1 and Der1, have retained the overall architecture of rhomboid proteases, they may also bind and unwind substrates; ultimately lowering the energetic barrier for substrate removal.

Work by Greenblatt et al. has elucidated much of the mechanistic actions of human derlin, Derlin-1, in ERAD retrotranslocation. Specifically, Derlin-1—just like its yeast paralog, Dfm1— requires the widely conserved rhomboid motifs, WR and GxxxG, for retrotranslocation (Greenblatt et al., 2011). Derlin-1 has been implicated in several diseases such as viral infection, cancer, cystic fibrosis and neurological dysfunctions (Kandel and Neal, 2020). Accordingly, determining the mechanistic features associated with human derlin function is critical for understanding its vast roles in normal physiology and pathology. We have discovered that retrotranslocation-deficient Dfm1 mutants are located at sites that are conserved in human derlin Derlin-1. Site-directed mutagenesis of Derlin-1 at these conserved sites ablates retrotranslocation of its multi-spanning substrate, CFTR-ΔF508. Furthermore, Dfm1’s mechanistic actions can be extended to human derlins with Derlin-1 employing both lipid thinning function along with substrate binding through its L1 domain. A recent cryo-EM study showed that Derlin-1 forms a tetrameric channel(Rao et al., 2021). The authors proposed a mechanism of a single transmembrane helix traversing through the Derlin-1 channel during retrotranslocation. Notably, **TM2-TM5** lines the inner channel of the tetramer complex. Based on our simulations, this is the region where the lipid bilayer is distorted significantly. Having significant lipid thinning within the interior core where substrate TM helices are predicted to transverse is a reasonable location for lowering the energetic barrier to aid in the retrotranslocation process. Moreover, **TM1-Loop1-TM3 gate** lines the outer channel of the core and is aligned with our data for making the first contact with incoming membrane substrates. The authors predicted many hydrophobic residues within Loop1 (including Derlin-1 I56 residue we characterized in this study for substrate binding) are required for substrate access through this gate. This suggests an intriguing model in which derlin-mediated retrotranslocation of membrane substrates utilize both rhomboid features and channel activity. A high-resolution structure of derlins with their respective substrate along with crosslinking experiments to map out how substrates bind to Derlin-1 would be invaluable for investigating this phenomenon in the future.

In yeast, it appears that yeast rhomboid-like proteins Dfm1 and Der1 are separate retrotranslocation factors for membrane and lumenal substrates, respectively. In contrast, mammalian derlins have evolved to retrotranslocate both lumenal and membrane proteins (Greenblatt et al., 2011; Huang et al., 2013; Lopez-Serra et al., 2014). It would be interesting in the future to understand how Derlin-1 along with its paralogs, Derlin-2 and Derlin-3, retrotranslocate a wide range of ERAD substrates. Furthermore, it will be a fruitful avenue to expand our understanding of Derlin-2 and Derlin-3 and determine whether they require similar sequence features as yeast Dfm1 and human Derlin-1 for retrotranslocation. In one sense, we can recapitulate this transition between compartmentalized retrotranslocation to the more general mammalian functions by selecting our still-non-functional Der1-WR-GxxxG mutants for variants that have acquired the ability to function like Dfm1(Fig.1).

Our study has sought out to understand the mechanisms associated with the widely critical function of multi-spanning membrane substrate retrotranslocation. The homology model of Dfm1 does not appear to resemble a channel despite its ability to facilitate in the extraction of multi-spanning membrane proteins. In contrast to this view, a recent structure of human Derlin-1 demonstrates it forms a complex, which is conducive to having channel activity. Similarly, Der1 along with E3 ligase Hrd1 form a channel to function in ERAD-L retrotranslocation. This implies that derlin rhomboid pseudoproteases may employ both channel activity and its rhomboid features through L1-mediated substrate binding and lipid thinning to aid in the dislocation of membrane and luminal substrates respectively. Overall, this study provides functional insights of derlin rhomboid pseudoproteases, which will ultimately aid in the therapeutic design against these rhomboid-like proteins that are associated with a plethora of maladies including cancer, cystic fibrosis, and neurological dysfunctions.

## AUTHOR CONTRIBUTIONS

S.E.N., A.N., N.W., A.C., S.G., N.S., T.K., and R.E.A. designed research; S.E.N., A.N., N.W., A.C., S.G., N.S., T.K., S.B., and R.K., performed research; S.E.N., A.N., N.W., A.C., S.G., N.S., T.K., and R.E.A. analyzed data; S.E.N., A.N., N.W., A.C., S.G., and R.E.A. wrote the paper. All authors reviewed the results and approved the final version of the manuscript.

## ACKNOWLEDGEMENTS

We thank Tom Rapoport (Harvard Medical School), Davis Ng (National University of Singapore), Randy Schekman (University of California, Berkeley), Susan Michaelis (John Hopkins University), Mark Hochstrasser (Yale School of Medicine), Jeff Brodsky (University of Pittsburgh) and Hideki Nishitoh (University of Miyazaki) for providing plasmids, antibodies and human cell lines. We also thank Dr. Randolph Hampton and Neal lab members for in depth discussions and technical assistance. These studies were supported by NIH grant 1R35GM133565-01, Burroughs Wellcome Fund 1013987 and Pew Biomedical Award (to S.E.N.). Simulations were run on the PSC Bridges system at the Pittsburgh Supercomputer Center through the XSEDE allocation NSF TG-CHE060063. S.E.N wishes to dedicate this work to her colleague, Sarah Holland. You will be missed.

## DECLARATION OF INTERESTS

The authors declare they have no competing interests within the contents of this article.

---

**Supplemental Figure 1.**
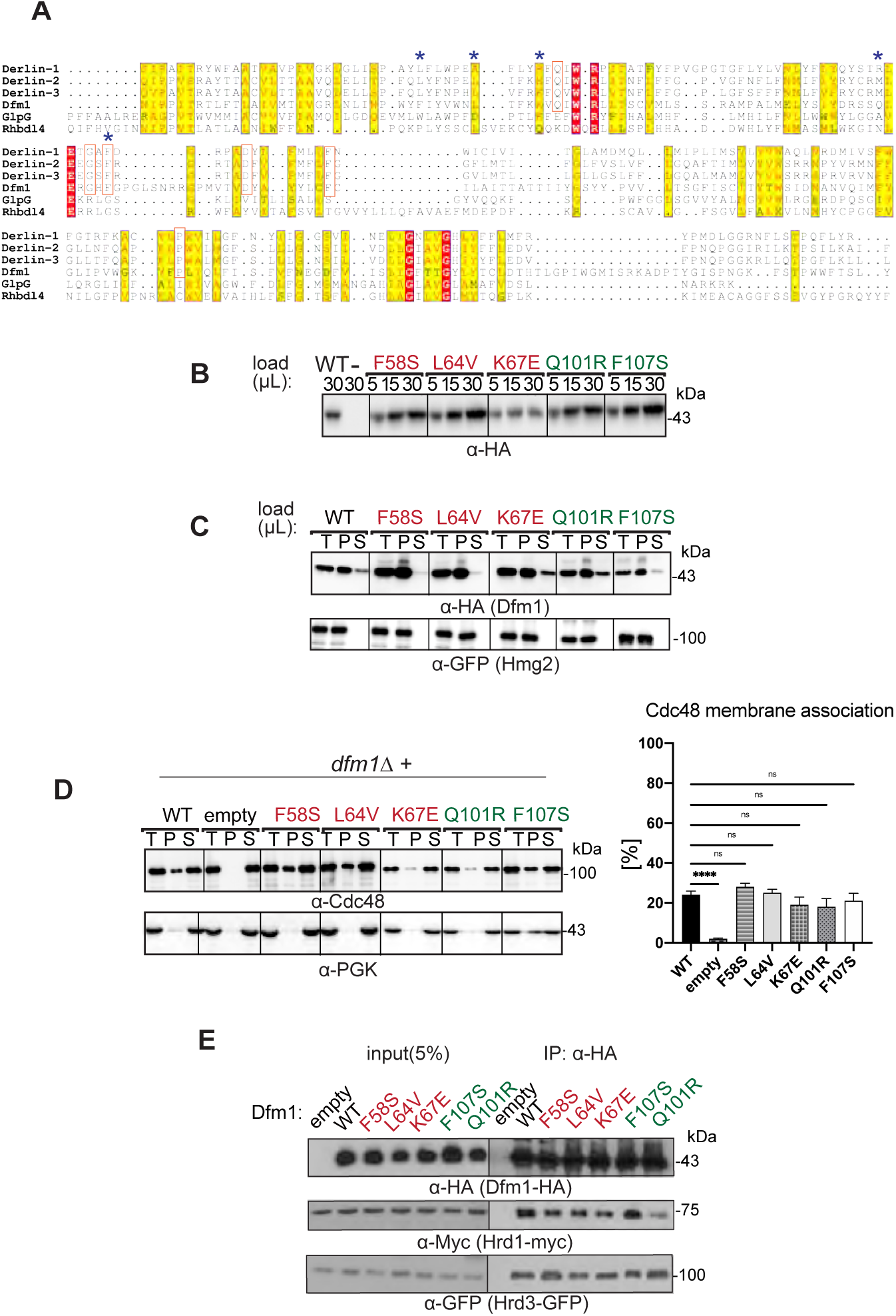
Related to Figure 1 &2. **(A)** Dfm1 is rhomboid pseudoprotease. TCoffee alignment of the transmembrane regions of Derlin-1, Derlin-2 and Derlin-3 and Rhbdl4 from H. sapiens; Dfm1 from S. cerevisiae; and Glpg from E. coli. Identically and similarly conserved residues are highlighted in red and yellow respectively. Furthermore, residues selected from loss of function screen is indicated by blue asterisks. **(B)** Dfm1 mutants are stably expressed. Stability of Dfm1 mutants were measured by loading increasing amounts of lysates (5 μL, 15 μL, and 30 μL) on SDS-PAGE followed by immunoblotting with α-HA. **(C)** Dfm1 mutants localize to the ER. Total cell lysate (T) from the indicated strains were separated into soluble cytosolic fraction (S) and pellet microsomal fraction (P) upon centrifugation at 14,000 x g. Each fraction was analyzed by SDS-PAGE and immunoblotted for Dfm1 mutants with α-HA and ER-localized Hmg2 with α-GFP. **(D)** Dfm1 mutants do not disrupt its Cdc48 recruitment function. Same as (A), except Cdc48 recruitment was analyzed by immunoblotting for Cdc48 with α-Cdc48 and Pgk1 with α-Pgk1. The graph shows the quantification of Cdc48 in the pellet fractions of the respective cells as measured from ImageJ. Data is represented as percentage of Cdc48 that is bound to pellet fraction and is shown as mean ± SEM from three independent experiments, **** p< 0.0001, Oneway ANOVA. **(E)** Association of retrotranslocation-deficient mutants to E3 ligase Hrd1 and Hrd3 was analyzed by co-IP. As a negative control, cells not expressing Dfm1 were used.

**Supplemental Figure 2.**
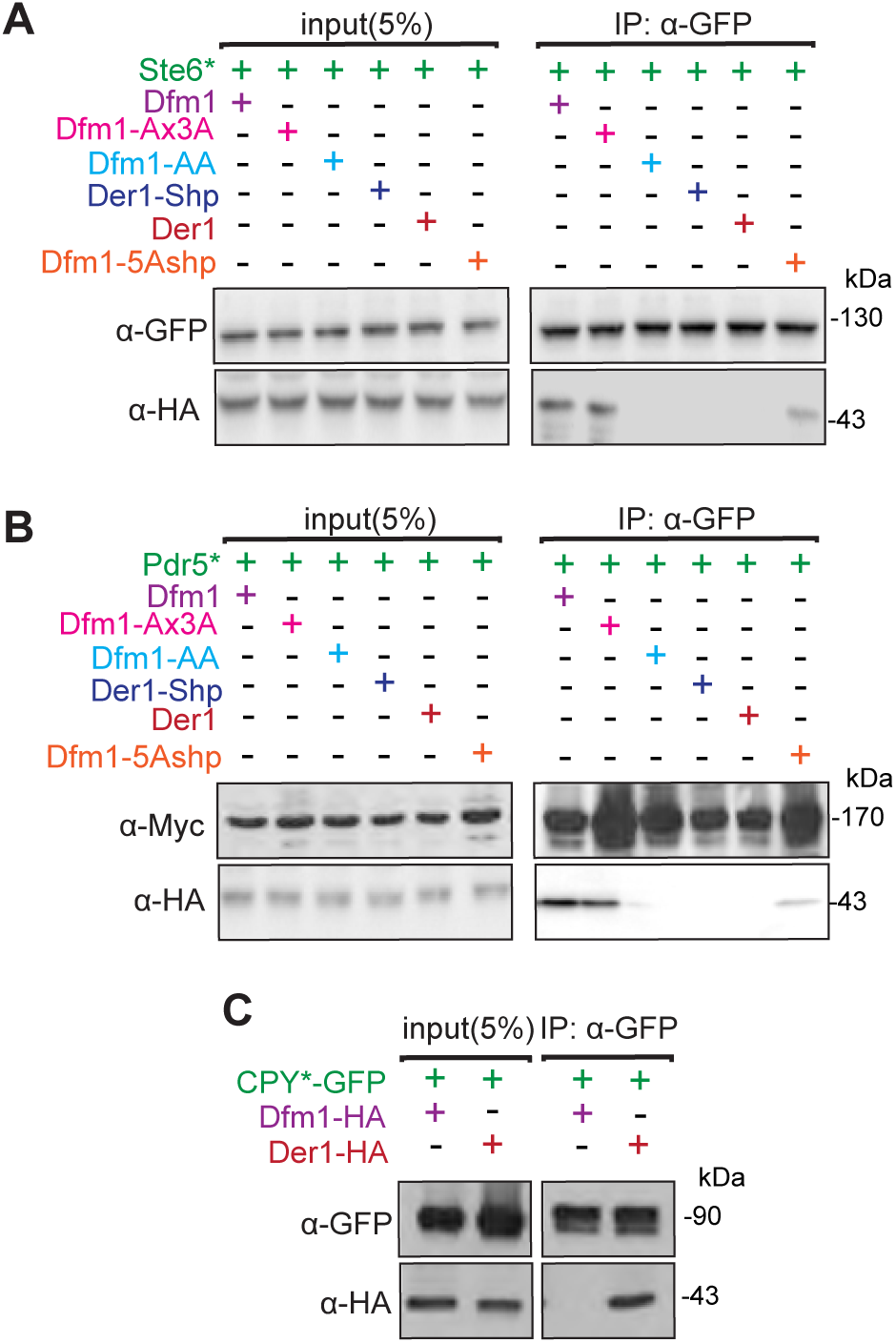
Related to Figure 4. **(A)** Co-IP was used to analyze Ste6* binding to Dfm1 variants; Dfm1-5Ashp, Der1-Shp, Dfm1-AA, and Dfm1-Ax3A. D was analyzed by co-IP. As a control for specificity, cells expressing Der1-HA was used. **(B)** Same as (A), except substrate Pdr5* was used in co-IP. **(C)** Same as (A) except substrate CPY* was used in co-IP.

**Supplemental Figure 3.**
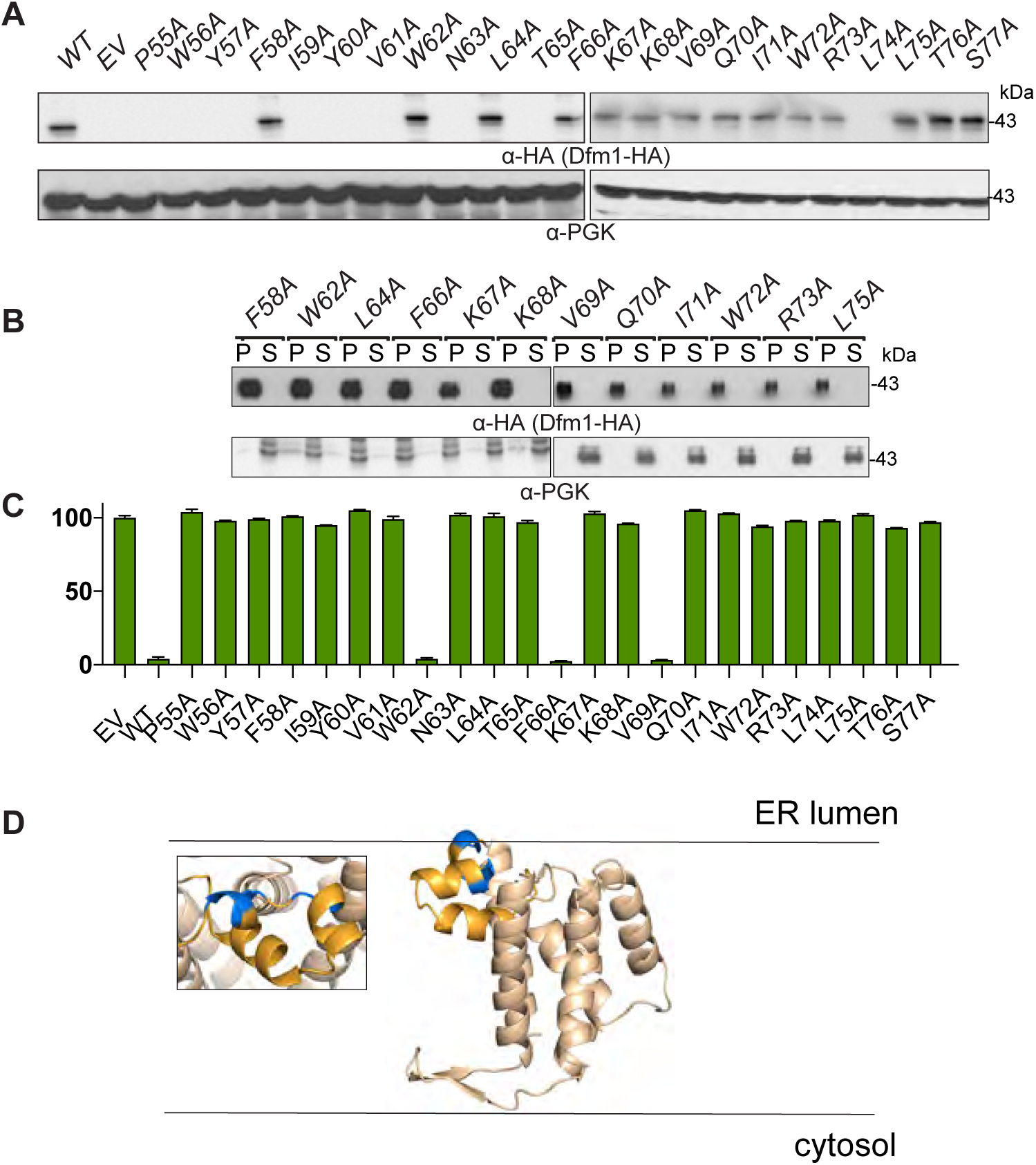
Related to Figure 5. **(A)** Stability of Dfm1 L1 mutants generated by Ala mutant scanning were measured by loading 30 μL of lysates on SDS-PAGE followed by immunoblotting with α-HA. **(B)** The indicated strains were grown to log phase and were subjected to flow cytometry to measure steady-levels of fluorescent membrane substrate, SUS-GFP. Histograms of 10,000 cells are shown, indicating the mean FITC-A value. **(C)** Dfm1 L1 mutants localize to the ER. The indicated strains were separated into soluble cytosolic fraction (S) and pellet microsomal fraction (P) upon centrifugation at 14,000 x g. Each fraction was analyzed by SDS-PAGE and immunoblotted for Dfm1 mutants with α-HA and cytosol-localized Pgk with α-Pgk. **(D)** Homology model of Dfm1. Position of non-polar residues (hydrophobic patch) are indicated in blue.

**Supplemental Figure 4.**
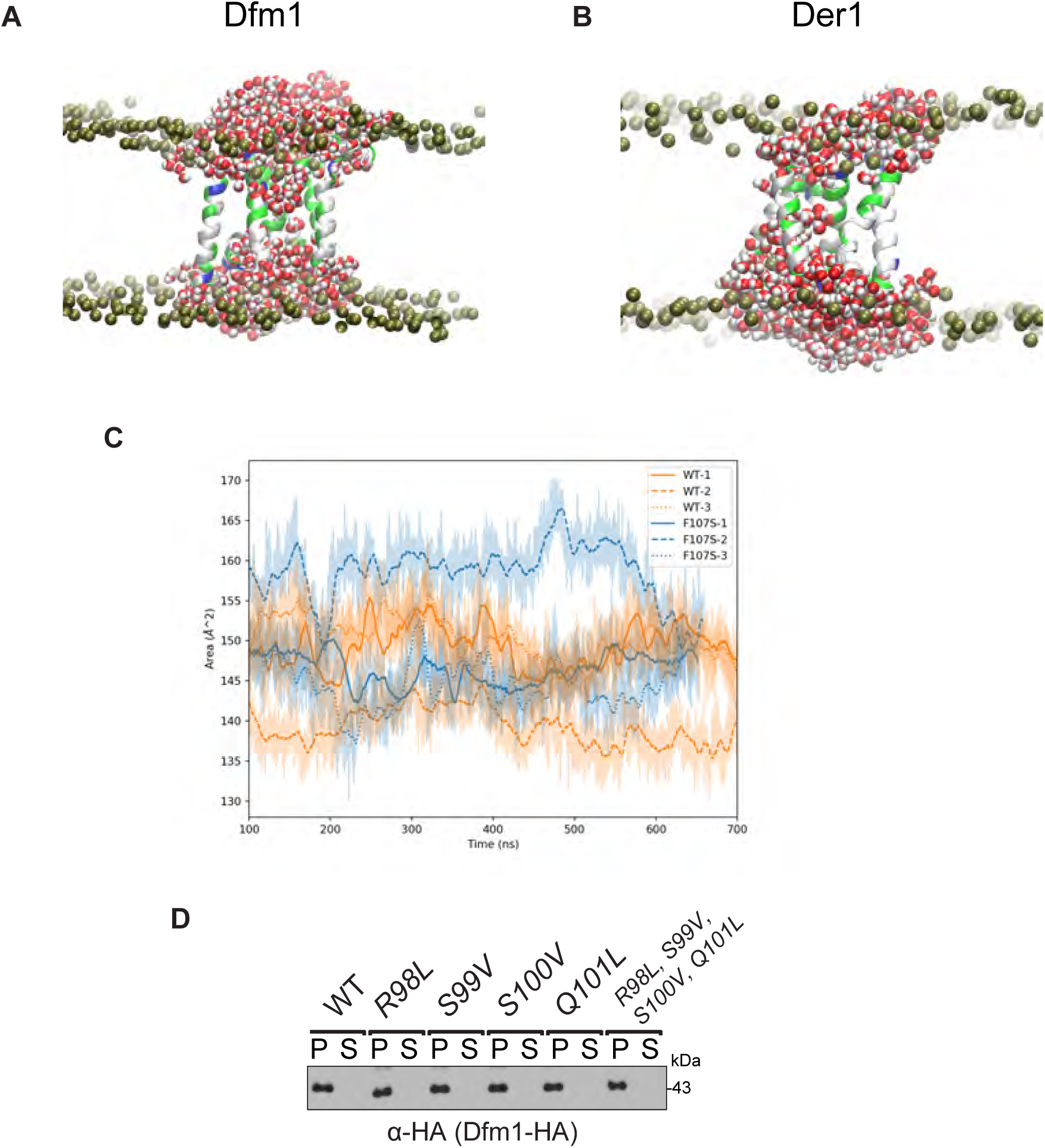
Related to Figure 6. **(A)** Simulation of S. cerevisiae derlin, Dfm1, homology model embedded in a mixed lipid bilayer (lipid composition in methods). Dfm1 is shown in multicolored ribbon representing the residue type (white is hydrophobic, green is polar, blue is positive, red is negative), water molecules are shown in red and white, and the phospholipid head group is shown in gold. **(B)** Same as (A), except the simulation was with S. cerevisiae derlin, Der1. **(C)** SASA calculation of the total exposed surface of WT and F107S Dfm1. The shaded area shows the raw data while the line marks a 10 ns moving average of the data.

**Supplemental Figure 5.**
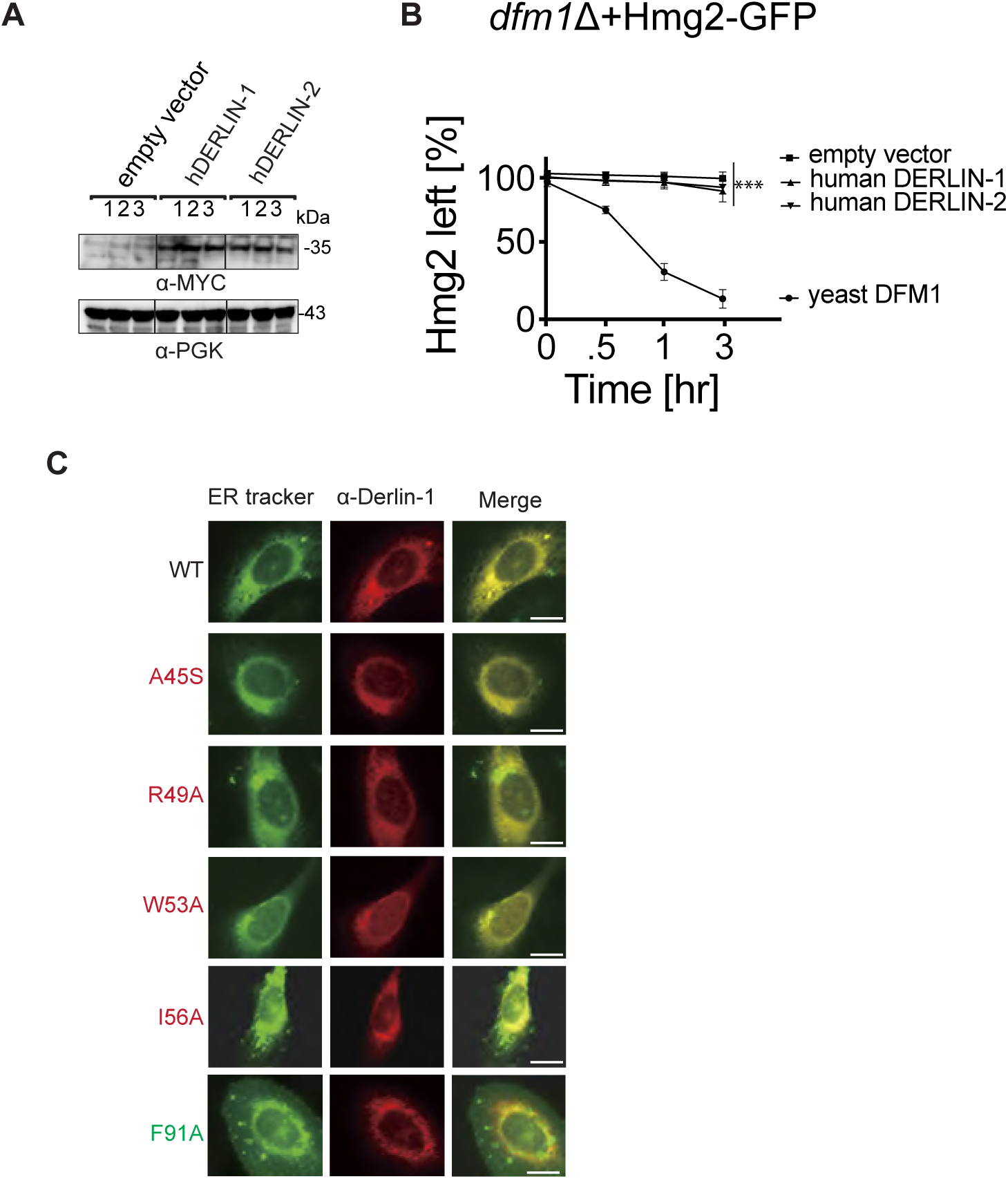
Related to Figure 7. **(A)** Indicated strains (n=3) heterogously expressing human Derlin-1 or Derlin-1 were grown to log-phase, lysed and analyzed by SDS-PAGE and immunoblotted for their steady-state elvels by alpha-Myc. (B) *dfm1*Δ strains containing Hmg2-GFP along with human Derlin-1, Derlin-2 or empty vector add back were grown to log-phase and degradation was measured by CHX-chase analysis. After CHX addition, Hmg2-GFP levels were measured by flow cytometry. Data is represented as mean ± SEM from three experiments, ***p < 0.001, Repeated Measures ANOVA. (C) HEK293T cells expressing indicated Derlin-1 mutants were stained with anti-MYC (Derlin-1, red) or DIOC6 (ER, green) and examined byconfocal microscopy in the mid-plane of each cell. Scale bar indicates 10 μM.

## STARS METHODS

### RESOURCE AVAILABILITY

#### Lead contact

Further information and requests for resources and reagents should be directed to and will be fulfilled by the Lead Contact, Sonya Neal.

#### Materials availability

Plasmids and yeast strains generated in this study is available from our laboratory.

#### Data and Code Availability

Original/source data for figures in the paper is available on Mendeley Data DOI.

### EXPERIMENTAL MODELS

All experiments were carried out in *Saccharomyces cerevisiae* budding yeast in BY4741 and S288C background and HEK293T cells (ATCC).

Authentication testing will be performed on established human cell lines regardless of the application, and testing will be done, at minimum, at the beginning and end of experimental work. For HEK293T human cell lines, short tandem repeat (STR) profiling will be performed and compared to results from online databases of human cell line STR profiles (ANSI/ATCC ASN-0002-2011 Authentication of Human Cell Lines: Standardization of STR Profiling. <u>ANSIeStandard Store</u>.)

### METHOD DETAILS

#### Yeast and Bacteria Growth Media

Standard yeast *Saccharomyces cerevisiae* growth media were used as previously described (Hampton and Rine, 1994), including yeast extract-peptone-dextrose (YPD) medium and ammonia-based synthetic complete dextrose (SC) and ammonia-based synthetic minimal dextrose (SD) medium supplemented with 2% dextrose and amino acids to enable growth of auxotrophic strains at 30°C. *Escherichia coli* Top10 cells were grown in standard LB media with ampicillin at 37°C as previously described (Gardner et al., 1998). HEK293 cells were cultured in DMEM medium supplemented with 10% FBS.

#### Plasmids and Strains

Plasmids used in this study are listed in Table S1. Plasmids for this work were generated using standard molecular biological techniques (Sato et al., 2009) and verified by sequencing (Eton Bioscience, Inc.). Primer information is available upon request. Full-length human DERLIN-1 cDNA was obtained by G-block synthesis (Eton Bioscience, Inc.) and subcloned into pcDNA3.1/Myc-His(+)A (Invitrogen) to express Derlin-1 with the myc epitope at the C-terminus. The KHN (pRH1958) and KWW (pRH1960) plasmids were a gift from Davis Ng (National University of Singapore, Singapore). The Ste6* plasmid (pRH2058) was a gift from S. Michaelis (Johns Hopkins School of Medicine, MD). The Pdr5* plasmid was a gift from Dieter H. Wolf (University of Stuttgart, Stuttgart, Germany). The pcDNA3.1-ΔF508-CFTR plasmid was a gift from J. Brodsky (University of Pittsburgh, PA).

A complete list of yeast strains and their corresponding genotypes are listed in Table S2. All strains used in this work were derived from S288C or Resgen. Yeast strains were transformed with DNA or PCR fragments using the standard LiOAc method (Ito et al., 1983). Null alleles were generated by using PCR to amplify a selection marker flanked by 50 base pairs of the 5’ and 3’ regions, which are immediately adjacent to the coding region of the gene to be deleted. The selectable markers used for making null alleles were genes encoding resistance to G418 or CloNat/nourseothricin. After transformation, strains with drug markers were plated onto YPD followed by replica-plating onto YPD plates containing (500 μg/mL G418 or 200 μg/mL nourseothricin). All gene deletions were confirmed by PCR.

#### *dfm1*Δ strain handling

To observe the phenotypic effect of *dfm1*Δ null strains, freshly transformed *dfm1*Δ null cells with the respective ERAD-M substrates was used in every assay.

#### Homology modeling

To build the 3D model of yeast derlin, Dfm1 protein on the template of yeast derlin Der1, the Phyre2 system was utilized(Kelley et al., 2015). Initially, the primary sequence is scanned against a database of 10 million known sequences for homologs via PSI-Blast. From here, homologous sequences are organized into an evolutionary fingerprint through Hidden Markov Models. Evolutionary fingerprints and Hidden Markov Models are made for the 65,000 known 3D structures to create a database of known structures. A scan of the evolutionary fingerprint with the database creates an alignment to known structures ranked by confidence of homology. This alignment generates a 3D threaded model with excellent accuracy even when sequence identity is less than 15%, and in addition is able to reliably detect extremely remote homology(Kelley et al., 2015).

#### Molecular dynamics simulation

The Dfm1 protein structure used in the MD simulations was built through homology modeling with the SWISS-MODEL structure prediction server. Der1 was the primary homologous structure for the predicted model of Dfm1 and the generated model covered residues 31-236 of Dfm1. Systems were prepared for MD using CHARMM-GUI’s membrane builder to place the protein in a 100Å X 100Å lipid patch and to apply any necessary amino acid mutations. The lipid composition was built to be representative of the ER membrane with a number percent composition of 47% POPC [1-palmitoyl-2-oleoyl-glycero-3-phosphocholine], 20% POPE [1-palmitoyl-2-oleoyl-sn-glycero-3-phosphoethanolamine], 15% cholesterol, 11% POPI [1-palmitoyl-2-oleoyl-sn-glycero-3-phosphoinositol], and 7% POPS [1-palmitoyl-2-oleoyl-sn-glycero-3-phospho-L-serine]. The system was solvated with the TIP3P water model and included 0.15M NaCl salt concentration. MD simulations were run with the CHARMM36m forcefield in an NPT ensemble at 310K and 1.01325 bar using the GROMACS 2018.3 MD engine. Systems were energy minimized and subsequently equilibrated in a stepwise manner, slowly relaxing the restraints, for a total of 2 ns, with the first 100ns of zero restraints simulation being considered as additional equilibration time. Both the wildtype protein and the F107S mutant were simulated in triplicate with each replicate running for ∼700ns. A set of simulations for Der1 (Fig S4) were run under the same conditions but for only ∼400 ns.

#### MD analyses

Membrane thickness calculations were performed using mdtraj to first split the lipids into separate leaflets and then calculate the distances between lipid headgroups of opposing leaflets for each frame of the simulation. Membrane thickness was defined to be an average of the 3 shortest trans-leaflet distances between headgroups for each lipid. Protein structures were clustered using GROMACS’ cluster command which conducted GROMOS clustering on the backbone atoms of the protein structure. SASA calculations were performed using the SASA command in GROMACS with a probe radius of 1.4Å. Visualizations were rendered using VMD and POVRay3.0.

#### Random mutagenesis of Dfm1

pRH2013 plasmid containing DFM1 driven from its native promoter was amplified by PCR using a high fidelity Phusion polymerase (control) and error prone Mutazyme 2 to introduce point mutations into DFM1. Specifically, 500ng of template DNA (pRH2013) and 20 cycles of PCR were used to obtain an average of 1-3 point mutations within DFM1, excluding the genetic region encoding the SHP motifs as per protocol instructions. Mutagenized DFM1 was amplified using high fidelity Phusion polymerase and treated with Dpn1 at 37°C overnight to digest the original unmutagenized template followed by PCR cleanup of mutagenized DFM1 using Promega wizard PCR cleanup kit. In parallel, backbone plasmid from pRH2013 was prepared by overnight digestion with Spe1 and PshA1 and then purified from 0.8% agarose gel. For homologous recombination of mutagenized DFM1 with pRH2013 backbone, linearized pRH2013 and purified mutagenized DFM1 were co-transformed into *dfm1Δhrd1Δ* yeast cells containing TDH3pr-SUS-GFP using a 1:9 backbone to insert ratio. Recombinants were selected on SC-Leu and incubated at 30°C. Resulting transformants were selected for high colony fluorescence, indicating their inability to degrade the optical retrotranslocation reporter, SUS-GFP. Plasmids were recovered from selected yeast transformants and transformed into *E. coli*.

Plasmids were recovered using Promega Wizard Plus SV Miniprep kit, as per manufacturer’s protocol and sent to ETON for sequencing using T7 (forward) and T3 (reverse) universal primers. Results for sequencing were aligned to wildtype DFM1 and mutated regions were identified using in house python scripts. Mutants containing one point mutation and no early stop codons verified by both forward and reverse strands were selected as mutants of interest.

#### Plasmid recovery from transformants

Plasmid extractions was performed as described in (Flagg et al., 2019). Transformants were inoculated into 3-ml YPD and grown overnight. The following day, 1 ml of YPD was added to stationary phase cultures, which were then allowed to grow for an hour at 30°. The entire culture was then pelleted and resuspended in 250 μl of resuspension buffer from a Promega Wizard Plus SV Miniprep kit (A1460). Resuspended cells were lysed with beads for 5 min in a multi-vortexer. Lysed cells and supernatant were then collected by nesting the microcentrifuge tube into a 15-ml conical tube, piercing the 2-ml microcentrifuge tube with a needle, and spinning the nested tubes at 2,000 rpm for 2 min. Lysed cells and cell lysate were then thoroughly resuspended, and the remainder of the miniprep was carried out according to the manufacturer’s protocol.

#### Cell culture, transfections and immunoblotting

Both wildtype and DERLIN-1 knockout HEK293T cell lines (ATCC) (kindly provided by Dr. Hideki Nishitoh, University of Miyazaki) were cultured in Dulbecco’s modified Eagle medium (25 mM glucose, sodium pyruvate) (Invitrogen) and grown at 37°C and 5% CO_2_. The media was supplemented with 10% fetal bovine serum (FBS) (Atlanta Biological). Both pcDNA3.1-DERLIN-1 and pcDNA3.1-ΔF508-CFTR were co-transfected into HEK293T cells in a 1:1 ratio using Lipofectamine^®^ LTX (Invitrogen) according to manufacturer’s instructions. Forty-eight hours after transfection, cells were lysed in lysis buffer (50 mM HEPES pH 7.5, 150 mM NaCl, 1% NP40, 0.1% SDS and 0.5 % sodium deoxycholate, and 1 mM EDTA supplemented with protease inhibitors for 1 hour on ice. After a centrifugation at 21,000 x *g* for 10 min at 4°C, the protein concentration was measured using the BCA protein assay kit (Pierce) and resuspended in SDS sample buffer and subjected to immunoblotting analysis. Equal amounts of protein extracts (30 μg) were separated by SDS-PAGE, transferred on nitrocellulose membrane and immunoblotted for anti-GAPDH (Bio-Rad), anti-Derlin-1 (ABclonal, Inc.) and anti-CFTR (CFTR Antibodies Distribution Program).

#### *In Vivo* Retrotranslocation Assay

*in vivo* retrotranslocation assay was performed as described in (Neal et al., 2019). Cells in log phase (OD_600_ 0.2-0.3) were treated with MG132 (benzyloxycarbonyl-Leu-Leu-aldehyde, Sigma) at a final concentration of 25 μg/mL (25 mg/mL stock dissolved in DMSO) for 2 hours at 30°C and GGPP (Geranylgeranyl pyrophosphate ammonium salt, Sigma) at a final concentration of 11 μM for 1 hour at 30°C and 15 ODs of cells were pelleted. Cells were resuspended in H_2_0, centrifuged and lysed with the addition of 0.5 mM glass beads and 400 μL of XL buffer (1.2 M sorbitol, 5 mM EDTA, 0.1 M KH_2_PO_4_, final pH 7.5) with PIs, followed by vortexing in 1minute intervals for 6-8 min at 4°C. Lysates were combined and clarified by centrifugation at 2,500 g for 5 min. Clarified lysate was ultracentrifuged at 100,000 g for 15 min to separate pellet (P100) and supernatant fraction (S100). P100 pellet was resuspended in 200 μL SUME (1% SDS, 8 M Urea, 10 mM MOPS, pH 6.8, 10 mM EDTA) with PIs and 5 mM N-ethyl maleimide (NEM, Sigma) followed by addition of 600 μL immunoprecipitation buffer (IPB) with PIs and NEM. S100 supernatant was added directly to IPB with PIs and NEM. 15 μL of rabbit polyclonal anti-GFP antisera (C. Zuker, University of California, San Diego) was added to P100 and S100 fractions for immunoprecipitation (IP) of Hmg2-GFP. Samples were incubated on ice for 5 minutes, clarified at 14,000 g for 5 min and removed to a new eppendorf tube and incubated overnight at 4°C. 100 μL of equilibrated Protein A-Sepharose in IPB (50% w/v) (Amersham Biosciences) was added and incubated for 2 h at 4°C. Proteins A beads were washed twice with IPB and washed once more with IP wash buffer (50 mM NaCl, 10 mM Tris), aspirated to dryness, resuspended in 2x Urea sample buffer (8 M urea, 4% SDS, 1mM DTT, 125 mM Tris, pH 6.8), and incubated at 55°C for 10 min. IPs were resolved by 8% SDS-PAGE, transferred to nitrocellulose, and immunoblotted with monoclonal anti-ubiquitin (Fred Hutchinson Cancer Center, Seattle) and anti-GFP (Clontech, Mountain View, CA). Goat anti-mouse (Jackson ImmunoResearch, West Grove, PA) and goat anti-rabbit (Bio-Rad) conjugated with horseradish peroxidase (HRP) recognized the primary antibodies. Western Lightning® Plus (Perkin Elmer, Watham, MA) chemiluminescence reagents were used for immunodetection.

#### Cycloheximide-Chase Assay

For yeast cells, cycloheximide chase assays were performed as previously described (Sato et al., 2009). Cells were grown to log-phase (OD_600_ 0.2-.03) and cycloheximide was added to a final concentration of 50 μg/mL. At each time point, a constant volume of culture was removed and lysed. Lysis was initiated with addition of 100 μl SUME with PIs and glass beads, followed by vortexing for 4 min. 100 μl of 2xUSB was added followed by incubation at 55°C for 10 min. Samples were clarified by centrifugation and analyzed by SDS-PAGE and immunoblotting. For HEK293T, cells were transfected with the indicated plasmids and 48 h later, cells were digested with trypsin and passaged in fresh medium supplemented with cycloheximide (20 μg/mL) for the indicated times. Equivalent volume of cell suspensions was harvested at different time points. Cell extracts were subjected to immunoblotting analysis as described above.

#### Cdc48 Microsome Association Assay

Yeast strains were grown to log phase (OD_600_ 0.2-0.3) and 15 ODs of cells were pelleted. Cells were resuspended in H_2_0, centrifuged and lysed with the addition of 0.5 mM glass beads and 400 μL of XL buffer with PIs and vortexed in 1-minute intervals for 6-8 min at 4°C. Lysates were combined and clarified by centrifugation at 2,500 g *for 5 min. 50 μL of lysate was transferred to another tube and designated as total fraction (T). The rest of* clarified lysate was centrifuged at 20,000 x g for 5 min to separate microsome pellet (P) and cytosolic supernatant fraction (S). An equivalent volume of 2xUSB was added to T, P and S fractions followed by solubilization at 55°C for 10 min. Samples were clarified by centrifugation, analyzed by SDS-PAGE and immunoblotted for Cdc48 and PGK1 with α-CDC48 (1:5,000) and α-PGK1(1:5,000) respectively.

#### Native Co-IP

Cultures from various yeast strains were grown to OD_600_ .2-.45 and 15 ODs of cells were pelleted, rinsed with H_2_0 and lysed with 0.5 mM glass beads in 400 μL of MF buffer supplemented with protease inhibitors. This was followed by vortexing at 1-minute intervals for 6-8 minutes at 4°C. Lysates were combined and clarified by centrifugation at 2,500 g for 5 min followed by centrifugation at 14,000 g for 15 min to obtain the microsomal pellet. The microsomal pellet was resuspended in 1 mL of Tween IP buffer (500 mM NaCl, 50 mM Tris, pH 7.5, 10 mM EDTA. 1.5% Tween-20) and incubated on ice for 30 minutes. Lysates were then centrifuged for 30 min at 14,000 x g, and the supernatant was incubated overnight with 10 μL of equilibrated GFP-Trap® agarose (ChromoTek Inc., Hauppauge, NY) at 4°C. The next day, the GFP-Trap® agarose beads were combined to one tube, washed once with non-detergent IP buffer, washed once more with IP wash buffer and resuspended in 100 μL of 2xUSB. Samples were resolved on 8% SDS-PAGE and immunoblotted for ubiquitin with anti-Ub, Cdc48 with α-CDC48, Hmg2-GFP with α-GFP, Dfm1-HA with α-HA, and Ste6*-GFP with α-GFP.

HEK293 cells were treated with 50 μM MG132 for 12 hours and lysed by douncing in minimal buffer (10 mM HEPES pH 8.0, 1 mM MgCl2, 2 mM KCl supplemented with protease inhibitor cocktails, and centrifuged at 35,000 rpm for 30 min at 4°C. Microsomal pellet was resuspended in solubilization buffer (50 mM HEPES pH 8.0, 300 mM NaCl, 10 mM imidizole, 1 mM MgCl_2_, 2 mM of KCl, 1% digitonin, and protease inhibitors) followed by incubation for 3 hours at 4°C. Lysates were centrifuged at 80,000 rpm for 30 min at 4°C followed by incubation of supernatant with Ni^+2^ beads at 4°C overnight. The next day, beads were washed twice with solubilization buffer, and incubated in elution buffer (solubilization buffer + 300 mM imidazole) to elute Derlin from Ni^+2^ beads, and resuspended in SDS sample buffer for immunoblotting analysis.

#### In Vivo Cross-linking

15 OD of cells from indicated strains were lysed with zymolyase and treated with vehicle (DMSO) or DSP (dithiobis(succinimidyl propionate) for 40 min. Microsomes were isolated and solubilized in Tween IP buffer supplemented with protease inhibitors and Hmg2-GFP was immunoprecipitated with GFP-Trap® agarose beads.

#### Free Polyubiquitin Competition Test

The ability of polyubiquitin to compete for Cdc48 binding to ubiquitinated Hmg2-GFP was adapted from Co-IP of Hmg2-GFP as described above. Microsomes were prepared and resuspended in 1 mL of Tween IP buffer supplemented with protease inhibitors followed by incubation on ice for 20 minutes. Lysates were then centrifuged for 30 min at 14,000 x g, and the supernatant was incubated with (2, 5, 10 or 20 μg) Lys48-linked polyubiquitin chains (BostonBiochem®) or buffer for one hour at 4°C. 30 μL of equilibrated GFP-Trap® agarose was added to each tube, and they were nutated overnight at 4°C. The next day, the GFP-Trap® agarose beads were washed with Tween IP buffer, washed once more with IP wash buffer and resuspended in 100 μL of 2xUSB. Samples were resolved on 8% SDS-PAGE and immunoblotted for Cdc48 with α-CDC48, Hmg2-GFP with α-GFP, and Dfm1-HA with α-HA.

#### Proteolytic Removal of Ubiquitin from Hmg2-GFP

Ubiquitin removal was accomplished with the broadly active Usp2 ubiquitin protease as previously described (Garza et al., 2009b), except that human recombinant Usp2Core (LifeSensors Inc., Malvern, PA) was used, and leupeptin and NEM were excluded from all buffers. Briefly, microsome fraction solubilized in 1 mL of Tween buffer was was incubated with 10 *μ*L of Usp2Core (5 *μ*g) for 1 hr at 37°C. The reaction was quenched with 200 *μl of* SUME with PIs and retrotranslocated Hmg2-GFP was immunoprecipitated as described above. 20 *μ*L of IP was used for detection of Hmg2-GFP with anti-GFP. 30 μL of equilibrated GFP-Trap® agarose was added and the sample was nutated overnight at 4°C. The next day, the GFP-Trap® agarose beads were washed with Tween IP buffer, washed once more with IP wash buffer and resuspended in 100 μL of 2xUSB. Samples were resolved on 8% SDS-PAGE and immunoblotted for ubiquitin with anti-Ub, Cdc48 with α-CDC48, Hmg2-GFP with α-GFP, and Dfm1-HA with α-HA.

#### Immunofluorescent staining

HEK293T cells were cultured on confocal slides, transfected with plasmids to express wildtype and Derlin-1 mutants. 36 hours later, cells were fixed with 2% paraformaldehyde at room temperature for 10 minutes, permeabilized with .1% Triton-X 100 in PBS, washed three times in 0.5% bovine serum albumin and 0.15% glycine at (pH 7.4) in phosphate-buffered saline, reacted with anti-His antibody (ABclonal), and incubated with Alexa 568 conjugated anti-mouse secondary antibody (ThermoFisher). After washing with 0.5% bovine serum albumin and 0.15% glycine at (pH 7.4) in phosphate-buffered saline, the coverslips were mounted for confocal microscopy.

### QUANTIFICATION AND STATISTICAL ANALYSIS

ImageJ (NIH) was used for all western blot quantifications. Band intensities were measured directly from films scanned in high resolution (600 dpi) in TIF file format. “Mean gray value” was set for band intensity measurements. In such experiments, a representative western blot was shown and band intensities were normalized to PGK1 loading control and quantified. t=0 was taken as 100% and data is represented as mean ± SEM from at least three experiments. GraphPad Prism was used for statistical analysis. Nested t-test, unpaired t-test or one-way factorial ANOVA followed by Bonferroni’s post-hoc analysis were applied to compare data. Significance was indicated as follow: n.s, not significant; * p<0.05, ** p<0.01, *** p<0.001, **** p<0.0001. The investigators were blinded during data analysis.

### SUPPLEMENTAL ITEMS

**Video Figure 1. Related to Figure 6**. Animated 2D contour plot of the membrane thickness every 10 ns of the simulation. The contour is plotted with the lines and colors binning thicknesses of every 0.2 nm between 1.0 and 5.0 nm.

**Table S1.**
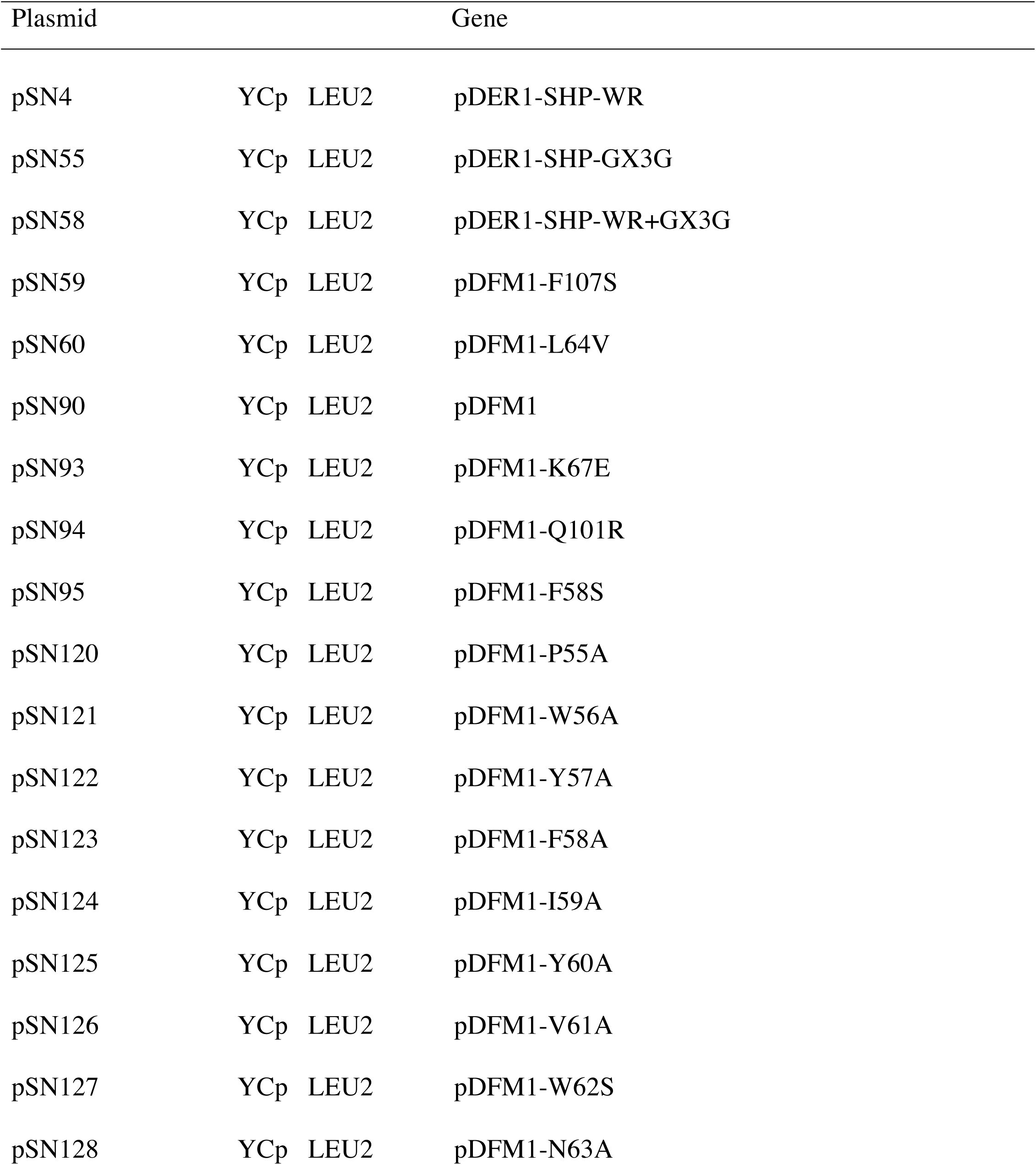
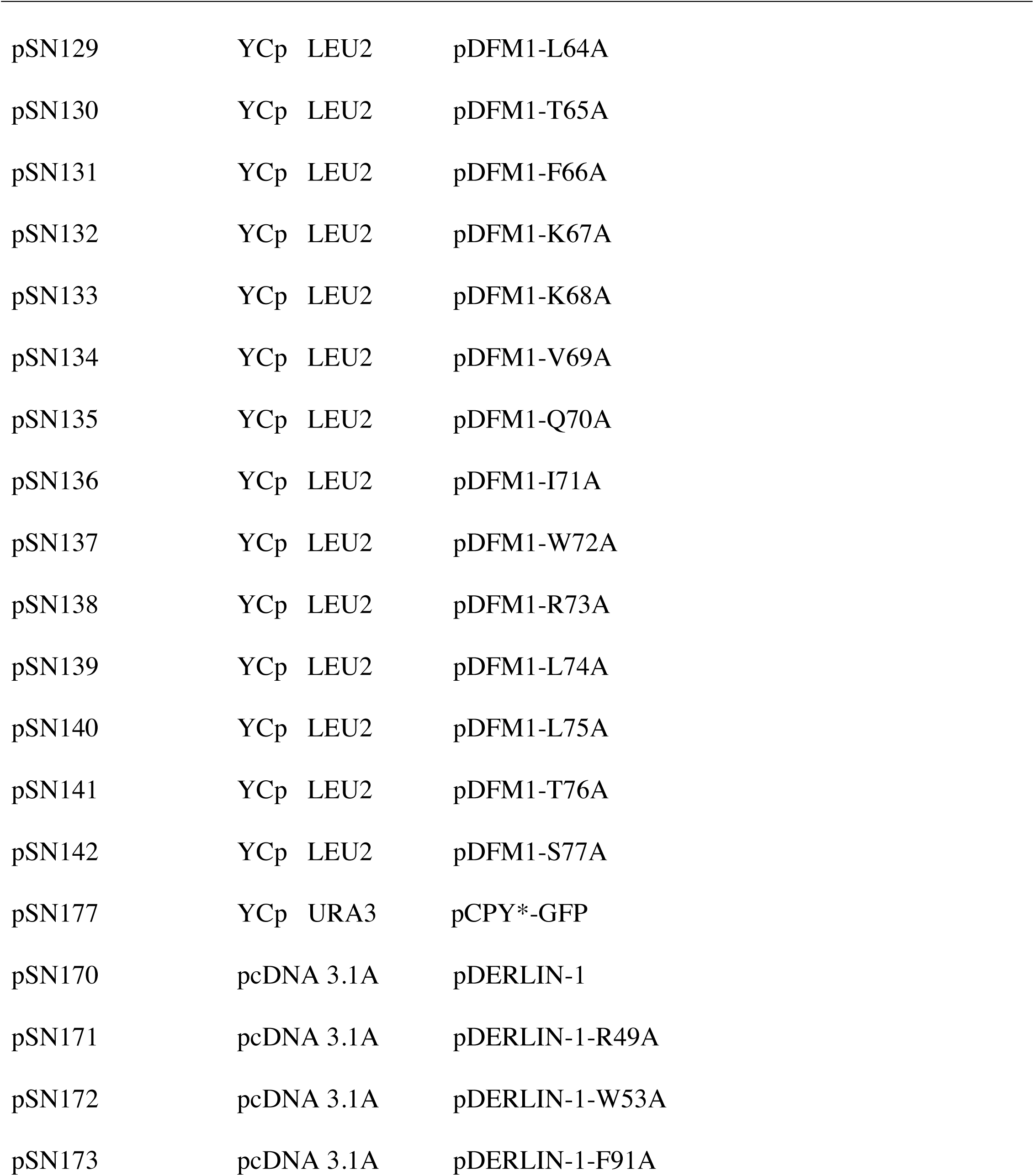
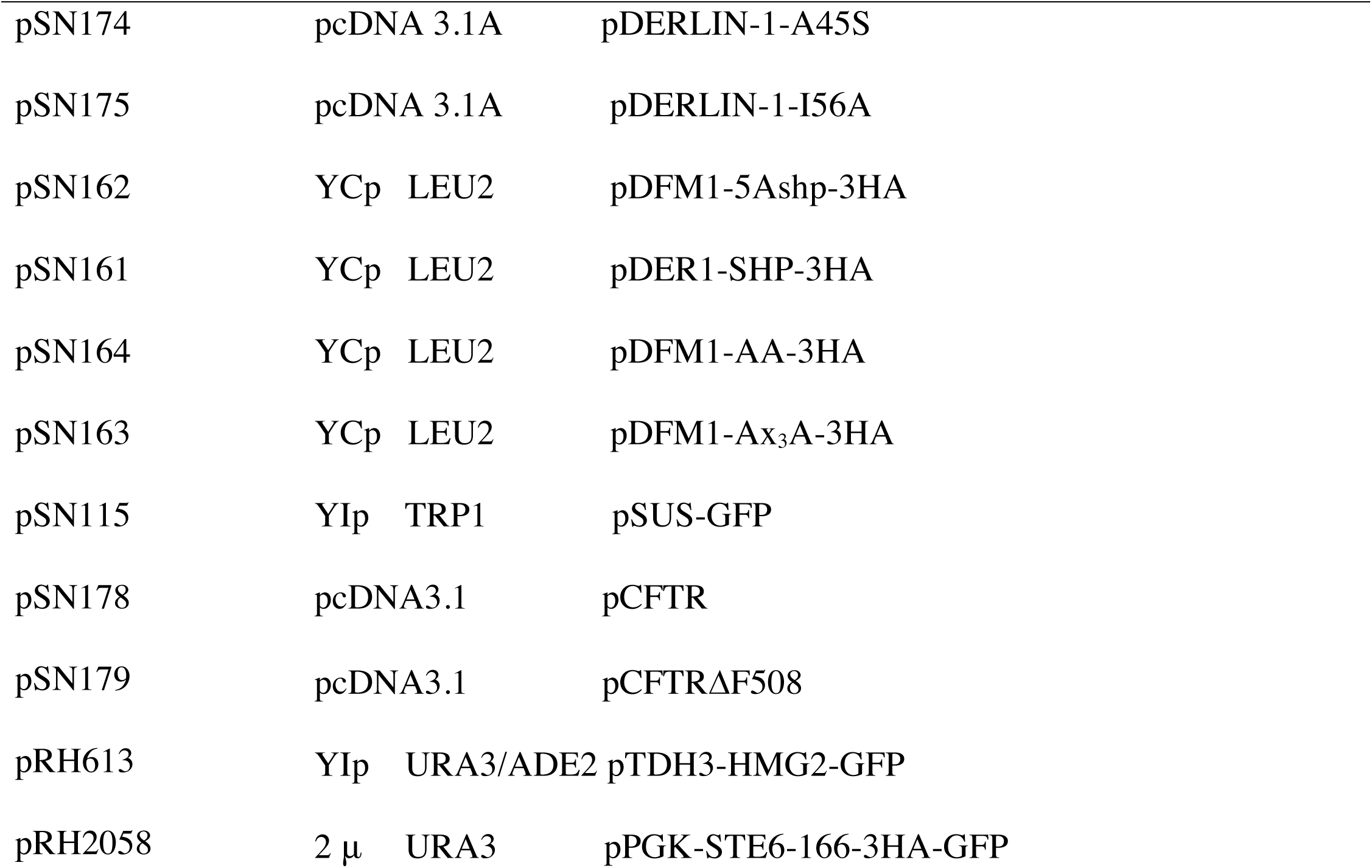
Plasmids used in this study, Related to Figures 1-7.

| Plasmid |  |  | Gene |
| --- | --- | --- | --- |
| pSN4 | YCp | LEU2 | pDER1-SHP-WR |
| pSN55 | YCp | LEU2 | pDER1-SHP-GX3G |
| pSN58 | YCp | LEU2 | pDER1-SHP-WR+GX3G |
| pSN59 | YCp | LEU2 | pDFM1-F107S |
| pSN60 | YCp | LEU2 | pDFM1-L64V |
| pSN90 | YCp | LEU2 | pDFM1 |
| pSN93 | YCp | LEU2 | pDFM1-K67E |
| pSN94 | YCp | LEU2 | pDFM1-Q101R |
| pSN95 | YCp | LEU2 | pDFM1-F58S |
| pSN120 | YCp | LEU2 | pDFM1-P55A |
| pSN121 | YCp | LEU2 | pDFM1-W56A |
| pSN122 | YCp | LEU2 | pDFM1-Y57A |
| pSN123 | YCp | LEU2 | pDFM1-F58A |
| pSN124 | YCp | LEU2 | pDFM1-I59A |
| pSN125 | YCp | LEU2 | pDFM1-Y60A |
| pSN126 | YCp | LEU2 | pDFM1-V61A |
| pSN127 | YCp | LEU2 | pDFM1-W62S |
| pSN128 | YCp | LEU2 | pDFM1-N63A |

|  |  |  |
| --- | --- | --- |
| pSN129 | YCp LEU2 | pDFM1-L64A |
| pSN130 | YCp LEU2 | pDFM1-T65A |
| pSN131 | YCp LEU2 | pDFM1-F66A |
| pSN132 | YCp LEU2 | pDFM1-K67A |
| pSN133 | YCp LEU2 | pDFM1-K68A |
| pSN134 | YCp LEU2 | pDFM1-V69A |
| pSN135 | YCp LEU2 | pDFM1-Q70A |
| pSN136 | YCp LEU2 | pDFM1-I71A |
| pSN137 | YCp LEU2 | pDFM1-W72A |
| pSN138 | YCp LEU2 | pDFM1-R73A |
| pSN139 | YCp LEU2 | pDFM1-L74A |
| pSN140 | YCp LEU2 | pDFM1-L75A |
| pSN141 | YCp LEU2 | pDFM1-T76A |
| pSN142 | YCp LEU2 | pDFM1-S77A |
| pSN177 | YCp URA3 | pCPY*-GFP |
| pSN170 | pcDNA 3.1A | pDERLIN-1 |
| pSN171 | pcDNA 3.1A | pDERLIN-1-R49A |
| pSN172 | pcDNA 3.1A | pDERLIN-1-W53A |
| pSN173 | pcDNA 3.1A | pDERLIN-1-F91A |

|  |  |  |
| --- | --- | --- |
| pSN174 | pcDNA 3.1A | pDERLIN-1-A45S |
| pSN175 | pcDNA 3.1A | pDERLIN-1-I56A |
| pSN162 | YCp LEU2 | pDFM1-5A<sub>shp</sub>-3HA |
| pSN161 | YCp LEU2 | pDER1-SHP-3HA |
| pSN164 | YCp LEU2 | pDFM1-AA-3HA |
| pSN163 | YCp LEU2 | pDFM1-A<sub>x3</sub>A-3HA |
| pSN115 | YIp TRP1 | pSUS-GFP |
| pSN178 | pcDNA3.1 | pCFTR |
| pSN179 | pcDNA3.1 | pCFTRΔF508 |
| pRH613 | YIp URA3/ADE2 | pTDH3-HMG2-GFP |
| pRH2058 | 2 $\mu$ URA3 | pPGK-STE6-166-3HA-GFP |

**Table S2.**
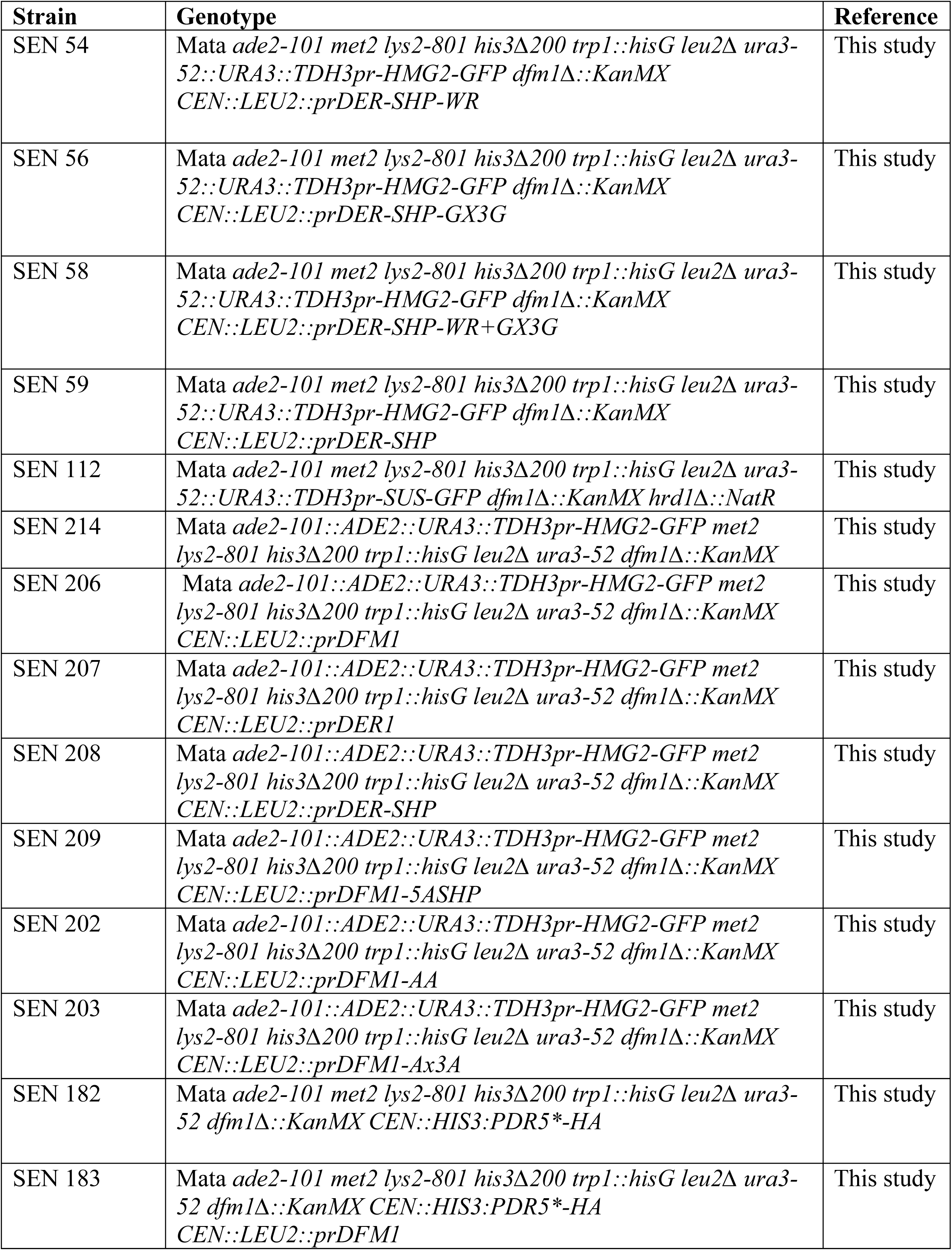
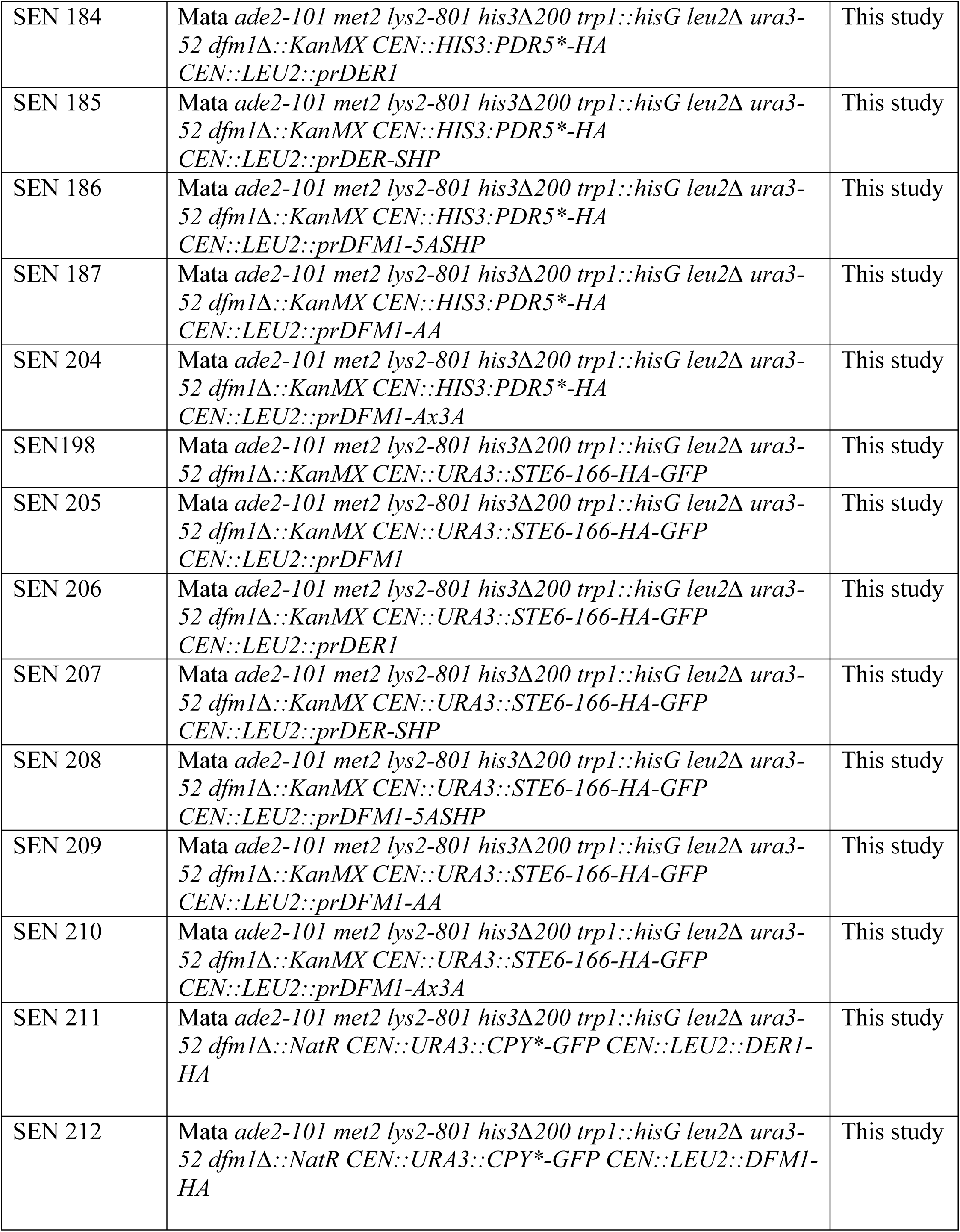
Yeast strains used in this study, Related to Figures 1-7.

**Table S3.** KEY RESOURCES TABLE. Related to Methods.

| REAGENT or RESOURCE | SOURCE | IDENTIFIER |
| --- | --- | --- |
| **Antibodies** |  |  |
| Mouse monoclonal anti-GFP | Clontech Laboratories, Inc. | Cat#632381; RRID: AB 2313808 |
| Mouse monoclonal anti-HA | Thermo Fisher Scientific | Cat#32-6700; RRID: AB 2533092 |
| Rabbit polyclonal anti-Myc | Genscript | Cat#A00172; RRID: AB 914457 |
| Rabbit polyclonal anti-Cdc48 | Neal et al., 2016 | N/A |
| Mouse monoclonal anti-PGK | Thermo Fisher Scientific | Cat#459250; RRID: AB 2569747 |
| Mouse monoclonal anti-Ubiquitin | Richard Gardner: University of Washington | N/A |
| Rabbit polyclonal anti-Derlin-1 | Abclonal | A8508 |
| Mouse anti-GAPDH |  |  |
| Rabbit anti-CFTR | Antibodies Distribution Program | N/A |
| **Bacterial and Virus Strains** |  |  |
| *Escherichia coli* Top10 Competent Cells | ThermoFisher Scientific | Cat#C404010 |
| **Biological Samples** |  |  |
| N/A |  |  |
| **Chemicals, Peptides, and Recombinant Proteins** |  |  |
| MG132 (benzyloxycarbonyl-Leu-Leu-aldehyde) | Sigma-Aldrich | 474787; CAS: 133407-82-6 |
| Cycloheximide | Sigma-Aldrich | C7698; CAS: 66-819 |
| Protein A Sepharose | GE Healthcare | 17-0780-01 |
| Usp2core | LifeSensors | DB501 |
| AQUApure Tetra-Ub Chains (K48-linked) | R&D Systems | UC-210B |
| DpnI restriction enzyme | New England Biolabs | R0176L |
| High Fidelity Phusion polymerase | New England Biolabs | M0530L |
| PCR Clean-Up System | Promega | A9282 |
| Ampicillin | Biopioneer | C0029 |
| Nourseothricin | Neta Scientific, Inc | RPI-N51200-1.0 |
| G418 | Biopioneer | C0050 |
| Lipofectamine® LTX (Invitrogen) | ThermoFisher Scientific | A12621 |
| GFP-Trap agarose | ChromoTek | gta-20 |
| Myc-Trap agarose | ChromoTek | yta-20 |
| Geranylgeranyl pyrophosphate ammonium salt | Millipore Sigma | G6025 |
| 3,3-Dithio- *bis*-(sulfosuccinimidyl)propionate (DSP) | Millipore Sigma | 322133 |
| Protein A-Sepharose | Millipore Sigma | GE17-0780-01 |
| DIOC6 (3,3'-Dihexyloxacarbocyanine Iodide) | ThermoFisher Scientific | D273 |
| **Critical Commercial Assays** |  |  |
| GeneMorph II Random Mutagenesis Kit | Agilent Technologies | 200550 |
| **Deposited Data** |  |  |
| N/A |  |  |
| **Experimental Models: Cell Lines** |  |  |
| Human: HEK293 cell line | Kadowaki et al., 2018 | ATCC |
| **Experimental Models: Organisms/Strains** |  |  |
| *Saccharomyces cerevisiae* BY4741 | GE Dharmacon | Cat#YSC1048 |
| *Saccharomyces cerevisiae* S288C | This study | N/A |
| Additional yeast strains used: refer to Table S2 | This study |  |
| **Oligonucleotides** |  |  |
| N/A |  |  |
| **Recombinant DNA** |  |  |
| Plasmids used: refer to Table S1 | This study |  |
| pRABBIT IgG IRES-EmGFP Positive Control Vecto | ThermoFisher Scientific | A39243 |
| **Software and Algorithms** |  |  |
| Prism 7 for Mac | GraphPad Software | https://www.graphpad.com/scientific-software/prism/ |
| Image J | NIH | https://imagej.nih.gov/ij/ |
| FlowJo | Vashistha et al., 2016 | https://www.flowjo.com/solutions/flowjo |
| BD Accuri C6 | BD Accuri | Cat # 653122 |
| PyMOL | Schrodinger, LLC | https://pymol.org/2/ |
| Protein Data Bank | RCSB PDB | www.rcsb.org |

## Notes

### Competing Interest Statement

The authors have declared no competing interest.

## REFERENCES

1. Avci, D., Fuchs, S., Schrul, B., Fukumori, A., Breker, M., Frumkin, I., Chen, C., Biniossek, M., Kremmer, E., Schilling, O., et al. (2014). The Yeast ER-Intramembrane Protease Ypf1 Refines Nutrient Sensing by Regulating Transporter Abundance. Mol. Cell 56, 630–640.

2. Began, J., Cordier, B., Březinová, J., Delisle, J., Hexnerová, R., Srb, P., Rampírová, P., Kožíšek, M., Baudet, M., Couté, Y., et al. (2020). Rhomboid intramembrane protease YqgP licenses bacterial membrane protein quality control as adaptor of FtsH AAA protease. EMBO J. e102935.

3. Bhattacharya, A., and Qi, L. (2019). ER-associated degradation in health and disease - From substrate to organism. J. Cell Sci. 132.

4. Bodnar, N.O., and Rapoport, T.A. (2017). Molecular Mechanism of Substrate Processing by the Cdc48 ATPase Complex. Cell 169, 722–735.e9.

5. Bondar, A.N. (2020). Phosphatidylglyerol Lipid Binding at the Active Site of an Intramembrane Protease. J. Membr. Biol. 253, 563–576.

6. Bondar, A.-N., del Val, C., and White, S.H. (2009a). Rhomboid Protease Dynamics and Lipid Interactions. Structure 17, 395–405.

7. Bondar, A.N., del Val, C., and White, S.H. (2009b). Rhomboid Protease Dynamics and Lipid Interactions. Structure 17, 395–405.

8. Brooks, C.L., and Lemieux, M.J. (2013). Untangling structure-function relationships in the rhomboid family of intramembrane proteases. Biochim. Biophys. Acta - Biomembr. 1828, 2862–2872.

9. Carvalho, P., Goder, V., and Rapoport, T.A. (2006). Distinct ubiquitin-ligase complexes define convergent pathways for the degradation of ER proteins. Cell 126, 361–373.

10. Chen, B., Mariano, J., Tsai, Y.C., Chan, A.H., Cohen, M., and Weissman, A.M. (2006). The activity of a human endoplasmic reticulum-associated degradation E3, gp78, requires its Cue domain, RING finger, and an E2-binding site. Proc. Natl. Acad. Sci. U. S. A. 103, 341–346.

11. Cho, J.A., Chinnapen, D.J.F., Aamar, E., te Welscher, Y.M., Lencer, W.I., and Massol, R. (2012). Insights on the trafficking and retro-translocation of glycosphingolipid-binding bacterial toxins. Front. Cell. Infect. Microbiol. 2, 51.

12. Düsterhöft, S., Künzel, U., and Freeman, M. (2017). Rhomboid proteases in human disease: Mechanisms and future prospects. Biochim. Biophys. Acta - Mol. Cell Res. 1864, 2200–2209.

13. Flagg, M.P., Kao, A., and Hampton, R.Y. (2019). Integrating after CEN Excision (ICE) Plasmids: Combining the ease of yeast recombination cloning with the stability of genomic integration. Yeast 36, 593–605.

14. Fleig, L., Bergbold, N., Sahasrabudhe, P., Geiger, B., Kaltak, L., and Lemberg, M.K. (2012). Ubiquitin-Dependent Intramembrane Rhomboid Protease Promotes ERAD of Membrane Proteins. Mol. Cell 47, 558–569.

15. Foresti, O., Ruggiano, A., Hannibal-Bach, H.K., Ejsing, C.S., and Carvalho, P. (2013). Sterol homeostasis requires regulated degradation of squalene monooxygenase by the ubiquitin ligase Doa10/Teb4. Elife 2, e00953.

16. Gardner, R., Cronin, S., Leader, B., Rine, J., Hampton, R., and Leder, B. (1998). Sequence determinants for regulated degradation of yeast 3-hydroxy-3-methylglutaryl-CoA reductase, an integral endoplasmic reticulum membrane protein. Mol. Biol. Cell 9, 2611–2626.

17. Garza, R.M., Sato, B.K., and Hampton, R.Y. (2009a). In vitro analysis of Hrd1p-mediated retrotranslocation of its multispanning membrane substrate 3-hydroxy-3-methylglutaryl (HMG)-CoA reductase. J. Biol. Chem. 284, 14710–14722.

18. Garza, R.M., Sato, B.K., and Hampton, R.Y. (2009b). In vitro analysis of Hrd1p-mediated retrotranslocation of its multSUSispanning membrane substrate 3-hydroxy-3-methylglutaryl (HMG)-CoA reductase. J. Biol. Chem. 284, 14710–14722.

19. Goder, V., Carvalho, P., and Rapoport, T. a (2008). The ER-associated degradation component Der1p and its homolog Dfm1p are contained in complexes with distinct cofactors of the ATPase Cdc48p. FEBS Lett. 582, 1575–1580.

20. Greenblatt, E.J., Olzmann, J. a, and Kopito, R.R. (2011). Derlin-1 is a rhomboid pseudoprotease required for the dislocation of mutant α-1 antitrypsin from the endoplasmic reticulum. Nat. Struct. Mol. Biol. 18, 1147–1152.

21. Greenblatt, E.J., Olzmann, J.A., and Kopito, R.R. (2012). Making the cut: intramembrane cleavage by a rhomboid protease promotes ERAD. Nat. Struct. Mol. Biol. 19, 979–981.

22. Ha, Y., Akiyama, Y., and Xue, Y. (2013). Structure and Mechanism of Rhomboid Protease. J. Biol. Chem. 288, 15430–15436.

23. Hampton, R.Y., and Garza, R.M. (2009). Protein quality control as a strategy for cellular regulation: lessons from ubiquitin-mediated regulation of the sterol pathway. Chem. Rev. 109, 1561–1574.

24. Hampton, R.Y., and Rine, J. (1994). Regulated degradation of HMG-CoA reductase, an integral membrane protein of the endoplasmic reticulum, in yeast. J. Cell Biol. 125, 299– 312.

25. Hampton, R.Y., and Sommer, T. (2012). Finding the will and the way of ERAD substrate retrotranslocation. Curr. Opin. Cell Biol. 24, 460–466.

26. Hampton, R.Y., Gardner, R.G., and Rine, J. (1996). Role of 26S proteasome and HRD genes in the degradation of 3-hydroxy-3-methylglutaryl-CoA reductase, an integral endoplasmic reticulum membrane protein. Mol. Biol. Cell 7, 2029–2044.

27. Huang, C.-H., Hsiao, H.-T., Chu, Y.-R., Ye, Y., and Chen, X. (2013). Derlin2 Protein Facilitates HRD1-mediated Retro-translocation of Sonic Hedgehog at the Endoplasmic Reticulum. J. Biol. Chem. 288, 25330–25339.

28. Hwang, J., Ribbens, D., Raychaudhuri, S., Cairns, L., Gu, H., Frost, A., Urban, S., and Espenshade, P.J. (2016). A Golgi rhomboid protease Rbd2 recruits Cdc48 to cleave yeast SREBP. EMBO J. 35, 2332–2349.

29. Ito, H., Fukuda, Y., Murata, K., and Kimura, A. (1983). Transformation of intact yeast cells treated with alkali cations. J. Bacteriol. 153, 163–168.

30. Kandel, R.R., and Neal, S.E. (2020). The role of rhomboid superfamily members in protein homeostasis: Mechanistic insight and physiological implications. Biochim. Biophys. Acta - Mol. Cell Res. 1867.

31. Kelley, L.A., Mezulis, S., Yates, C.M., Wass, M.N., and Sternberg, M.J.E. (2015). The Phyre2 web portal for protein modeling, prediction and analysis. Nat. Protoc. 10, 845– 858.

32. Kreutzberger, A.J.B., Ji, M., Aaron, J., Mihaljević, L., and Urban, S. (2019). Rhomboid distorts lipids to break the viscosity-imposed speed limit of membrane diffusion. Science (80-.). 363.

33. Laney, J.D., and Hochstrasser, M. (2003). Ubiquitin-dependent degradation of the yeast Matα2 repressor enables a switch in developmental state. Genes Dev. 17, 2259–2270.

34. Lemberg, M.K., and Adrain, C. (2016). Inactive rhomboid proteins: New mechanisms with implications in health and disease. Semin. Cell Dev. Biol. 60, 29–37.

35. Lemberg, M.K., and Freeman, M. (2007). Functional and evolutionary implications of enhanced genomic analysis of rhomboid intramembrane proteases. Genome Res. 17, 1634–1646.

36. Lemieux, M.J., Fischer, S.J., Cherney, M.M., Bateman, K.S., and James, M.N.G. (2007). The crystal structure of the rhomboid peptidase from Haemophilus influenzae provides insight into intramembrane porteolysis. Proc. Natl. Acad. Sci. U. S. A. 104, 750–754.

37. Lilley, B.N., and Ploegh, H.L. (2004). A membrane protein required for dislocation of misfolded proteins from the ER. Nature 429, 834–840.

38. Lopez-Serra, P., Marcilla, M., Villanueva, A., Ramos-Fernandez, A., Palau, A., Leal, L., Wahi, J.E., Setien-Baranda, F., Szczesna, K., Moutinho, C., et al. (2014). A DERL3-associated defect in the degradation of SLC2A1 mediates the Warburg effect. Nat. Commun. 5, 3608.

39. Marinko, J.T., Huang, H., Penn, W.D., Capra, J.A., Schlebach, J.P., and Sanders, C.R. (2019). Folding and Misfolding of Human Membrane Proteins in Health and Disease: From Single Molecules to Cellular Proteostasis. Chem. Rev. 119, 5537–5606.

40. Mehrtash, A.B., and Hochstrasser, M. (2019a). Ubiquitin-dependent protein degradation at the endoplasmic reticulum and nuclear envelope. Semin. Cell Dev. Biol. 93, 111–124.

41. Mehrtash, A.B., and Hochstrasser, M. (2019b). Ubiquitin-dependent protein degradation at the endoplasmic reticulum and nuclear envelope. Semin. Cell Dev. Biol. 93, 111–124.

42. Moin, S.M., and Urban, S. (2012). Membrane immersion allows rhomboid proteases to achieve specificity by reading transmembrane segment dynamics. Elife 2012.

43. Moon, H.W., Han, H.G., and Jeon, Y.J. (2018). Protein quality control in the endoplasmic reticulum and cancer. Int. J. Mol. Sci. 19.

44. Müller, D.J., Kessler, M., Oesterhelt, F., Möller, C., Oesterhelt, D., and Gaub, H. (2002). Stability of bacteriorhodopsin α-helices and loops analyzed by single-molecule force spectroscopy. Biophys. J. 83, 3578–3588.

45. Neal, S., Mak, R., Bennett, E.J., and Hampton, R. (2017). A Cdc48 “retrochaperone” function is required for the solubility of retrotranslocated, integral membrane Endoplasmic Reticulum-associated Degradation (ERAD-M) substrates. J. Biol. Chem. 292.

46. Neal, S., Jaeger, P.A., Duttke, S.H., Benner, C.K., Glass, C., Ideker, T., and Hampton, R. (2018). The Dfm1 Derlin Is Required for ERAD Retrotranslocation of Integral Membrane Proteins. Mol. Cell 69.

47. Neal, S., Duttke, S.H., and Hampton, R.Y. (2019). Assays for protein retrotranslocation in ERAD.

48. Neal, S., Syau, D., Nejatfard, A., Nadeau, S., and Hampton, R.Y. (2020). HRD Complex Self-Remodeling Enables a Novel Route of Membrane Protein Retrotranslocation. IScience 23.

49. Peterson, B.G., Glaser, M.L., Rapoport, T.A., and Baldridge, R.D. (2019). Cycles of autoubiquitination and deubiquitination regulate the erad ubiquitin ligase hrd1. Elife 8.

50. Plemper, R.K., Egner, R., Kuchler, K., and Wolf, D.H. (1998). Endoplasmic reticulum degradation of a mutated ATP-binding cassette transporter Pdr5 proceeds in a concerted action of Sec61 and the proteasome. J. Biol. Chem. 273, 32848–32856.

51. Rao, B., Li, S., Yao, D., Wang, Q., Xia, Y., Jia, Y., Shen, Y., and Cao, Y. (2021). The cryo-EM structure of an ERAD protein channel formed by tetrameric human Derlin-1. Sci. Adv. 7.

52. Ravid, T., Kreft, S.G., and Hochstrasser, M. (2006). Membrane and soluble substrates of the Doa10 ubiquitin ligase are degraded by distinct pathways. EMBO J. 25, 533–543.

53. Ruggiano, A., Foresti, O., and Carvalho, P. (2014). ER-associated degradation: Protein quality control and beyond. J. Cell Biol. 204, 869–879.

54. Sato, B.K., and Hampton, R.Y. (2006a). Yeast Derlin Dfm1 interacts with Cdc48 and functions in ER homeostasis. Yeast 1053–1064.

55. Sato, B.K., and Hampton, R.Y. (2006b). Yeast Derlin Dfm I interacts with Cdc48 and functions in ER homeostasis. Yeast 23, 1053–1064.

56. Sato, B.K., Schulz, D., Do, P.H., and Hampton, R.Y. (2009). Misfolded membrane proteins are specifically recognized by the transmembrane domain of the Hrd1p ubiquitin ligase. Mol. Cell 34, 212–222.

57. Schmidt, C.C., Vasic, V., and Stein, A. (2020). Doa10 is a membrane protein retrotranslocase in er-associated protein degradation. Elife 9, 1–31.

58. Shokhen, M., and Albeck, A. (2017). How does the exosite of rhomboid protease affect substrate processing and inhibition? Protein Sci. 26, 2355–2366.

59. Sicari, D., Igbaria, A., and Chevet, E. (2019). Control of Protein Homeostasis in the Early Secretory Pathway: Current Status and Challenges. Cells 8.

60. Stolz, A., Schweizer, R.S., Schäfer, A., and Wolf, D.H. (2010a). Dfm1 forms distinct complexes with Cdc48 and the ER ubiquitin ligases and is required for ERAD. Traffic 11, 1363–1369.

61. Stolz, A., Schweizer, R.S., Schäfer, A., and Wolf, D.H. (2010b). Dfm1 Forms Distinct Complexes with Cdc48 and the ER Ubiquitin Ligases and Is Required for ERAD. Traffic 11, 1363–1369.

62. Sun, Z., and Brodsky, J.L. (2019). Protein quality control in the secretory pathway. J. Cell Biol. 218, 3171–3187.

63. Sun, F., Zhang, R., Gong, X., Geng, X., Drain, P.F., and Frizzell, R.A. (2006a). Derlin-1 Promotes the Efficient Degradation of the Cystic Fibrosis Transmembrane Conductance Regulator (CFTR) and CFTR Folding Mutants. J. Biol. Chem. 281, 36856–36863.

64. Sun, F., Zhang, R., Gong, X., Geng, X., Drain, P.F., and Frizzell, R.A. (2006b). Derlin-1 promotes the efficient degradation of the cystic fibrosis transmembrane conductance regulator (CFTR) and CFTR folding mutants. J. Biol. Chem. 281, 36856–36863.

65. Suzuki, M., Otsuka, T., Ohsaki, Y., Cheng, J., Taniguchi, T., Hashimoto, H., Taniguchi, H., and Fujimoto, T. (2012). Derlin-1 and UBXD8 are engaged in dislocation and degradation of lipidated ApoB-100 at lipid droplets. Mol. Biol. Cell 23, 800–810.

66. Tichá, A., Collis, B., and Strisovsky, K. (2018). The Rhomboid Superfamily: Structural Mechanisms and Chemical Biology Opportunities. Trends Biochem. Sci. 43, 726–739.

67. Uritsky, N., Shokhen, M., and Albeck, A. (2016). Stepwise Versus Concerted Mechanisms in General-Base Catalysis by Serine Proteases. Angew. Chem. Int. Ed. Engl. 55, 1680–1684.

68. Vashist, S., and Ng, D.T.W. (2004). Misfolded proteins are sorted by a sequential checkpoint mechanism of ER quality control. J. Cell Biol. 165, 41–52.

69. Wang, J.-Z., and Dehesh, K. (2018). ER: the Silk Road of interorganellar communication. Curr. Opin. Plant Biol. 45, 171–177.

70. Wang, Y., Zhang, Y., and Ha, Y. (2006). Crystal structure of a rhomboid family intramembrane protease. Nature 444, 179–183.

71. Wang, Y., Maegawa, S., Akiyama, Y., and Ha, Y. (2007). The Role of L1 Loop in the Mechanism of Rhomboid Intramembrane Protease GlpG. J. Mol. Biol. 374, 1104–1113.

72. Wangeline, M.A., and Hampton, R.Y. (2018). “Mallostery”—ligand-dependent protein misfolding enables physiological regulation by ERAD. J. Biol. Chem. 293, 14937–14950.

73. Wu, X., Siggel, M., Ovchinnikov, S., Mi, W., Svetlov, V., Nudler, E., Liao, M., Hummer, G., and Rapoport, T.A. (2020). Structural basis of ER-associated protein degradation mediated by the Hrd1 ubiquitin ligase complex. Science (80-.). 368, 1–13.

74. Xiong, L., Zhang, L., Yang, Y., Li, N., Lai, W., Wang, F., Zhu, X., and Wang, T. (2020). ER complex proteins are required for rhodopsin biosynthesis and photoreceptor survival in Drosophila and mice. Cell Death Differ. 27, 646–661.

75. Yagishita, N., Yamasaki, S., Nishioka, K., and Nakajima, T. (2008). Synoviolin, protein folding and the maintenance of joint homeostasis. Nat. Clin. Pract. Rheumatol. 4, 91–97.

76. Ye, Y., Shibata, Y., Yun, C., Ron, D., and Rapoport, T.A. (2004). A membrane protein complex mediates retro-translocation from the ER lumen into the cytosol. Nature 429, 841–847.

77. You, H., Ge, Y., Zhang, J., Cao, Y., Xing, J., Su, D., Huang, Y., Li, M., Qu, S., Sun, F., et al. (2017). Derlin-1 promotes ubiquitylation and degradation of the epithelial Na+ channel, ENaC. J. Cell Sci. 130.

78. Zhou, Y., Moin, S.M., Urban, S., and Zhang, Y. (2012). An internal water-retention site in the rhomboid intramembrane protease GlpG ensures catalytic efficiency. Structure 20, 1255–1263.

79. Zoll, S., Stanchev, S., Began, J., Škerle, J., Lepšík, M., Peclinovská, L., Majer, P., and Strisovsky, K. (2014). Substrate binding and specificity of rhomboid intramembrane protease revealed by substrate–peptide complex structures. EMBO J. 33, 2408–2421.

